# Measuring expression heterogeneity of single-cell cytoskeletal protein complexes

**DOI:** 10.1101/2020.09.12.294801

**Authors:** Julea Vlassakis, Louise L. Hansen, Ryo Higuchi-Sanabria, Yun Zhou, C. Kimberly Tsui, Andrew Dillin, Haiyan Huang, Amy E. Herr

**Author notes:** These authors contributed equally (Julea Vlassakis, Louise L. Hansen). Corresponding Author: Amy E. Herr.

## Abstract

Multimeric cytoskeletal protein complexes orchestrate normal cellular function. However, protein-complex distributions in stressed, heterogeneous cell populations remain unknown. Cell staining and proximity-based methods have limited selectivity and/or sensitivity for endogenous multimeric protein-complex quantification from single cells. We introduce micro-arrayed, differential detergent fractionation to simultaneously detect protein complexes in 100s of individual cells. Fractionation occurs by 60 s size-exclusion electrophoresis with protein complex-stabilizing buffer that minimizes depolymerization. Co-detection of cytoskeletal protein complexes in U2OS cells treated with filamentous actin (F-actin) destabilizing LatA detects a subpopulation (~11%) exhibiting downregulated F-actin, but upregulated microtubule and intermediate filament protein complexes. Thus, some cells upregulate other cytoskeletal complexes to counteract the stress of LatA treatment. We also sought to understand the effect of non-chemical stress on cellular heterogeneity of F-actin. We find heat shock dysregulates F- and G-actin correlation. The assay overcomes selectivity limitations to biochemically quantify single-cell protein complexes perturbed with diverse stimuli.

## Introduction

Over 80,000 protein complexes comprised of interacting proteins regulate processes from proteostasis to transcription^1^. A critical set of multimeric protein complexes form the cell cytoskeleton: actin filaments, microtubules, and intermediate filaments. For example, actin dynamically polymerizes and depolymerizes^2,3^ between monomeric G-actin (~42 kDa) and filamentous F-actin^4^ to determine cell morphology, motility, and proliferation^5^. F-actin is considered the “functional” actin species in the cytoskeleton. Thus, the F-actin ratio (or F-actin abundance divided by total actin) is a metric for cytoskeletal integrity. F-actin levels can be increased in metastatic cancer cells^5^, thus underpinning the design of oncology drugs that disrupt F-actin filaments^6^. In addition, microtubule stabilizing chemotherapeutics (e.g., taxanes) are widely used in treatment of numerous cancers^7^ (e.g., breast, lung, prostate). However, development of taxane-resistant cell subpopulations^8^ requires further advances to screen drugs that target the cytoskeleton. Quantifying the distribution of cytoskeletal protein complexes in single cells would inform drug development and help elucidate stress-induced cancer transformations.

To understand cytoskeletal protein-complex expression heterogeneity, no existing method combines the needed detection sensitivity, throughput, and selectivity for multimeric protein complexes in single cells. Single-cell bottom-up mass spectrometry has been demonstrated,^9,10^ but cannot determine protein-complex stoichiometry like top-down mass spectrometry. However, lossy sample fractionation in top-down mass spectrometry prevents identification of protein complexes from low-cell number samples^11^. Targeted approaches such as proximity ligation assay and FRET achieve single-cell sensitivity, but rely on adjacent oligo-bound antibodies or fluorescent probes to infer that two proteins are interacting^12,13^. As proximity-based techniques are designed to measure 1:1 interaction, multi-component or multimeric protein complexes are difficult to detect. Finally, actin-specific detection methods are numerous, but suffer from limitations impacting sensitivity and selectivity and applicability to other cytoskeletal protein complexes. Visualization of the actin cytoskeleton relies on fluorescently tagged actin (e.g., GFP-actin fusion or split GFP-actin fusion^14^), GFP-fused actin binding proteins or peptides (e.g., Lifeact, F-tractin, Utrophin), nanobodies^15^, or chemicals that directly bind actin (e.g., phalloidin). Such molecules may alter cytoskeletal dynamics both *in vitro* and *in vivo*^16-18^. Phalloidin competes with, or is dissociated by, endogenous actin-binding proteins^19,20^ and actin-targeting drugs, such as Jasplakinolide^21^. Bulk ultracentrifugation overcomes these limitations while sacrificing single-cell resolution. In bulk ultracentrifugation, mild lysis in F-actin stabilization buffer solubilizes G-actin and preserves F-actin. The supernatant (G-actin) and pellet (F-actin) fractions are subsequently quantified by western blot or DNase inhibition assay^22^. However, bulk ultracentrifugation typically requires ~10^7^ cells, masking underlying cell-to-cell variation^22^.

To address gaps in multimeric protein-complex quantification, we introduce Single-cell protein Interaction Fractionation Through Electrophoresis and immunoassay Readout, or SIFTER. With sequential differential detergent fractionation and bi-directional, single-cell polyacrylamide gel electrophoresis (originally developed for nuclear versus cytoplasmic protein separation^23^), we electrophoretically separate monomers from protein complexes. Single cells are gravity-settled in microwells formed in polyacrylamide, where the microwell aspect ratio is selected to maximize singlecell microwell occupancy^24^. Here, each cell is lysed in situ in buffer designed to maintain protein complexes. Under an applied electric field, the gel size-excludes protein complexes larger than ~740 kDa to the microwell. Small monomeric proteins electromigrate into the gel in two steps: first from the monomeric fraction, and second from the intentionally depolymerized protein complex fraction after a buffer exchange. SIFTER fractionates protein complexes in < 1 min, or 40× faster than the ultracentrifugation assay. The thin SIFTER gel (0.5 mm thick) minimizes resistive heating that could prematurely depolymerize or dissociate protein complexes. Owing to the arrayed format and open microfluidic design, hundreds of fractionation separations are performed simultaneously. After fractionation and bi-directional electrophoresis, both the depolymerized protein complex (e.g., F-actin, microtubule, and/or intermediate filament) and monomer (e.g., G-actin, β-tubulin, or vimentin) states are blotted or immobilized in distinct gel regions abutting each microwell. Protein complex and monomer states are quantified by in-gel immunoprobing, allowing target multiplexing^24^. We applied SIFTER to four basic questions. First, do two well-studied actin-targeting drugs (Latrunculin A and Jasplakinolide) induce variation in F-actin complex-levels in single cells compared to controls? Second, as a corollary, does Latrunculin A yield cellular phenotypes distinct from controls with different organization of other cytoskeletal protein complexes, such as microtubules and intermediate filaments? Third, what is the distribution of the F-actin ratio across a population of single cells? Fourth, how does heat shock, another cellular stress, shift the F-actin ratio distribution and coordination between F- and G-actin at the singlecell level? We show SIFTER is a versatile method for understanding cellular heterogeneity – at singlecell resolution – in protein-complex levels in response to perturbation.

## Results

### SIFTER design principles and characterization

To selectively detect cytoskeletal protein complexes from single cells, we integrate differential detergent fractionation, electrophoretic separation, and immunoassay steps into a single microdevice. An important set of dynamic protein complexes comprise the cytoskeleton, including F-actin filaments, microtubules (MT) and intermediate filaments (IF; Fig. 1a). Two design considerations are central to our measurement of dynamic protein complexes: (1) discerning the protein complexes from monomers; and (2) maintaining protein complexes during fractionation. For the first design consideration, we focus on the F-actin filament, which is the smallest and most dynamic of the three cytoskeletal protein complexes. Each filament can be composed of up to 100s of globular G-actin monomers (k_off_ ~ 0.2 - 1.0 s^-1^ *in vivo*^25^). F-actin averages ~2.7 MDa (versus MT at ~178 MDa with 1 μm average MT length^26^ and 1625 tubulin heterodimers per μm of MT^27^, and IF at ~30 MDa for typical μm-scale IF^28^ at >30 kDa per nm of filament^29^). F-actin polymerization proceeds rapidly once four G-actin are incorporated in a filament. Steady-state polymerization (k_on_ ~0.1-5 μM^-1^s^-1^)^25^ yields a distribution of filament masses^30^. While the F-actin mass distribution below ~2700 kDa is unknown *in vivo*, F-actin is highly enmeshed. Thus, discerning F- (<160 kDa) vs. G-actin (42 kDa) requires coarse size cutoff (~100s of kDa), which should also fractionate MT and IF. On the second design consideration, rapid F-actin depolymerization occurs below the critical concentration of total actin (~0.2 - 2.0 μM *in vivo*). To maintain local concentrations of actin above the critical concentration demands < ~10× dilution during the assay, as cellular total actin is ~10-100 μM. Thus, the SIFTER fractionation gel contains microwells with ~10^8^× smaller reaction volume versus bulk ultracentrifugation to minimize dilution. The microwells accommodate gravity-sedimented single cells^24^ within the fractionation gel (Fig. 1a). The open SIFTER device is suited to rapid serial introduction of buffers via interchangeable hydrogel lids to first lyse cells and stabilize protein complexes during fractionation, and then depolymerize or dissociate protein complexes to spatially separate monomers from protein complexes (Fig. 1b-c).

**Figure 1:**
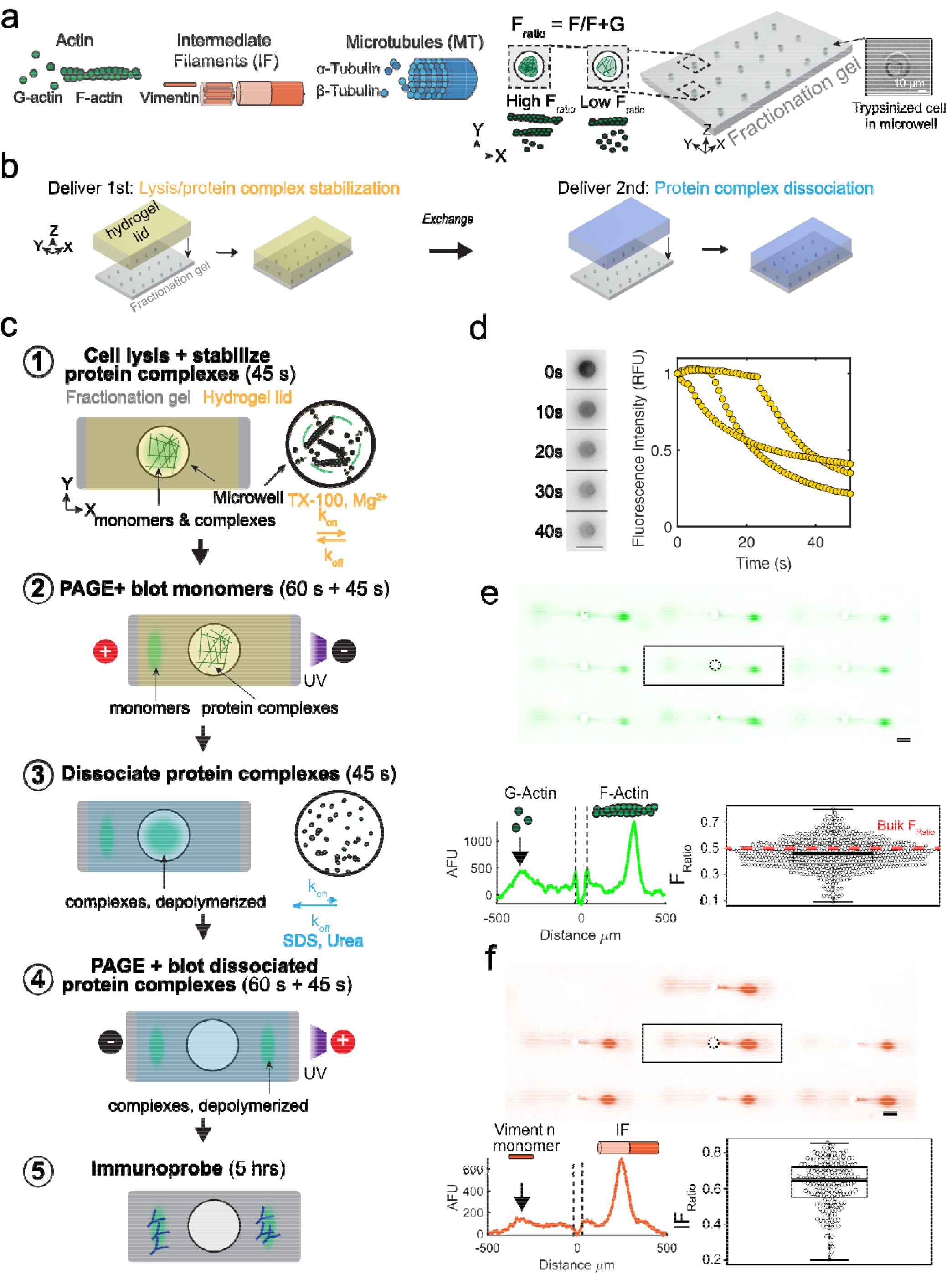
SIFTER detects cytoskeletal complexes from hundreds of single cells by on-chip integration of single-cell differential detergent fractionation and immunoblotting. (a) Schematic of three key cytoskeletal protein complexes: filamentous actin (F-actin; 4-100s of globular G-actin, 42 kDa each), microtubules (MT; assembled from alpha and β-tubulin heterodimers) and intermediate filaments (IF; comprising vimentin monomers). Trypsinized cells contain the three cytoskeletal protein complexes and are heterogeneous with low to high F-actin ratios (*F_ratio_*). Brightfield image shows a cell gravity settled into an individual microwell of a polyacrylamide fractionation gel. (b) Side-view schematic of hydrogel lid delivery of assay-stage optimized buffers to microwells in the fractionation gel. (c) The SIFTER assay comprises: 1) hydrogel lid delivery of protein complex-stabilizing lysis buffer to the array; 2) polyacrylamide gel electrophoresis (PAGE) and UV-immobilization of monomers (e.g., G-actin) in the gel; 3) hydrogel lid delivery of protein-complex dissociation buffer; 4) EP of dissociated protein complexes (e.g., F-actin, depolymerized) in the opposite direction of monomers and UV immobilization; and 5) in-gel antibody immunoprobing. (d) Cell lysis monitoring: false-color fluorescence micrograph montage and quantification of single MDA GFP-actin cells in microwells upon lysis with F-actin stabilization buffer (lyses cell but retains F-actin). Scale bar is 100 μm. Total fluorescence in the microwell normalized to initial in-microwell fluorescence as a function of lysis time for n=3 cells. (e) Immunoassay results: representative false-color micrograph of subset of the SIFTER array and intensity profile of GFP F-actin and GFP G-actin from single MDA-MB-231 GFP-actin cells (scale bar is 100 μm). Microwell annotated with dashed line. Boxplot with beeswarm shows the *F_ratio_* calculated from F- and G-actin peak area-under-the-curve from n = 692 single-cell protein complex separations from N = 3 SIFTER devices. Red dashed line shows the *F_ratio_* from a bulk assay^35^. (f) Representative false-color micrograph of a subset of the SIFTER array and intensity profile of vimentin from single MDA-MB-231 GFP-actin cells (scale bar is 100 μm). Boxplot with beeswarm shows *IF_ratio_* = *IF/VIM* from n = 215 cells from N = 4 SIFTER devices.

To report both the state (protein complex *vs.* monomer) and amount of specific protein complexes per cell, SIFTER comprises five assay steps (Fig. 1c). First, single trypsinized cells are gravity-settled in the microwell array (as described previously^31^) and lysed in an F-actin stabilization buffer delivered by the hydrogel lid, creating a lysate containing the monomers and complexes. Second, protein complexes are fractionated from the smaller monomers by polyacrylamide gel electrophoresis (PAGE, 60 s), during which large protein complexes are size-excluded from the gel and retained in each microwell. Monomers electrophorese into the gel and are immobilized (blotted) using a UV-induced covalent reaction to benzophenone methacrylamide integrated into the gel polymer network^24^. Third, to depolymerize the complexes retained in each microwell, a protein-complex depolymerization buffer is introduced by another hydrogel lid. Fourth, we electrophorese the now depolymerized complexes into a region of the gel separate from the immobilized monomers, where depolymerized complexes are in turn immobilized. Fifth, in-gel immunoprobing detects the immobilized populations of monomer and monomer depolymerized from the complexes (Fig. 1e-f) using a fluorescently labeled antibody probe against the protein (i.e., anti-actin antibody probe to detect F- and G-actin, and anti-vimentin antibody probe to detect vimentin monomers and intermediate filaments).

To maintain intact protein complexes in each microwell during PAGE fractionation, the F-actin stabilization buffer slows the natural depolymerization kinetics. The non-ionic detergent Triton X-100 at ~1% v/v lyses the cell and minimally alters in vitro polymerization rates of actin ^22, 32, 33^. The addition of 2 mM MgCl_2_ stabilizes F-actin complexes^22^, as Mg^2+^ binds G-actin to lower depolymerization rates^30^. Consequently, only ~2% of total F-actin depolymerizes per minute in mammalian cells lysed in stabilization buffer^22^, compatible with our goal to fractionate in ~1 min. Cell lysis depends on diffusion of Triton X-100 micelles, which requires ~10 s to reach the bottom of the microwells^34^. Imaging release of monomeric G-actin fused to fluorescent GFP from GFP-actin expressing breast cancer cells (MDA-MB-231 GFP-actin) within a microwell confirms a 45 s lysis yields only ~2.5-4× dilution of total actin to remain above the critical concentration (Fig. 1d). Important to minimizing F-actin-complex depolymerization during the assay, SIFTER completes cell lysis and fractionation in <5 min, or ~40× faster than bulk ultracentrifugation.

### Validation and benchmarking SIFTER

We first validated SIFTER by fractionating and quantifying the G-actin monomer *vs.* F-actin complexes in single MDA-MB-231 GFP-actin cells. We selected GFP-actin expressing cells to utilize fluorescence imaging to optimize cell lysis (Fig. 1d) and PAGE conditions. Immunoprobing for GFP yields distinct Gaussian protein peaks corresponding to GFP G-actin *(G)* on the left and GFP F-actin *(F)* to the right of each microwell (Fig. 1e). The area-under-the-curve of F-actin and G-actin peaks corresponds to the Factin (F) and G-actin (G) protein fraction abundances, respectively. We calculate the *F-actin ratio* (*F_ratio_*) = *F* / (*F*+*G*) for each cell. The MDA-MB-231 GFP-actin fusion cell average *F_ratio_* = 0.46 ± 0.11 (standard deviation; n = 692 cells, from N = 3 SIFTER devices), in reasonable agreement *with F_ratio_* ~0.5 for MDA-MB-231 from bulk ultracentrifugation^35^. With SIFTER, the *F_ratio_* coefficient of variation is 24%, revealing single-cell variation obscured in the bulk assay. *F_ratio_* variation measured by SIFTER includes cellular variation, such as the inverse correlation between the *F_ratio_* and cell volume. For example, cells grown in microniches that controllably decrease cell volume by half undergo similar magnitude increase in F-actin and decrease in G-actin (which should correspond to ~2× increase in *F_ratio_*)^36^.

Further, the F-actin stabilization buffer also maintains IF complexes (Fig. 1f, Supplementary Note 1). As such, we define and quantify an *IF ratio* (*IF_ratio_*) = *IF/VIM*, where *VIM* is the total amount of vimentin. The *IF_ratio_* indicates the fraction of vimentin actively giving structure to the cell, the primary function of IF. We find MDA-MB-231 GFP-actin cells have an average *IF_ratio_* = 0.62 ± 0.14 (error is the standard deviation; n = 215 cells, from N = 4 SIFTER devices measured on the same day). The significance of determining metrics such as *F_ratio_* and *IF_ratio_* with single-cell resolution is to detect small subpopulations of cells with distinctive filament and monomer distributions, especially the phenotypes that arise in response to stresses. Observed cell-to-cell variation in *F_ratio_* and *IF_ratio_* raises the intriguing question of whether cells compensate levels of one cytoskeletal protein complex for another. We investigate differential stress responses and compensation of cytoskeletal protein complexes later in this work.

To validate monomers *vs.* protein complex detection specificity, we determined the gel composition needed to fractionate F-actin (the smallest of the three cytoskeletal protein complexes) and directly observed PAGE of fluorescently labeled actin from single-cell lysates. The molecular mass cutoff for the gel depends on the total acrylamide concentration (%T). Based on native PAGE^37,38^, the SIFTER cutoff for an 8%T gel is ~740 kDa (Supplementary Fig. S1, Fig. 2a), or larger than 42 kDa G-actin, but smaller than an average ~2700 kDa F-actin. During PAGE of MDA-MB-231 GFP-actin cells (in which GFP is fused to both G- and F-actin), actin species indeed fractionate at the microwell edge (Fig. 2b). Within 45 s of PAGE, the G-actin Gaussian protein band completely injects 233 ± 11 μm into the polyacrylamide gel (with peak width sigma of 38.6 ± 5.3 μm, n=162; errors are standard deviations). We confirm the actin state of the species in the microwell by imaging PAGE of U2OS cells expressing RFP-Lifeact, a common marker for F-actin^16^. The microwell retains the F-actin complexes (Fig. 2c), with signal decrease attributable to diffusive losses^31^ of RFP-Lifeact-bound G-actin out of the microwell and photobleaching. We hypothesize two factors lead to no observed F-actin electromigration into the gel, including RFP-Lifeact bound dimers^39^. First, small oligomers are a minor fraction of F-actin due to substantial dissociation rates^40^. Second, highly crosslinked filaments^20^ remain enmeshed within the cytoskeleton even in lysed cells^41^. Further, we expect that free RFP-Lifeact would diffuse out of the microwell during cell lysis if present. Thus, we confirm that SIFTER fractionates F-actin complexes from single cells. Importantly, size exclusion may fractionate other protein complexes by adjusting the %T, as >99% of individual proteins of the mammalian proteome are larger than the molecular mass cutoff of even a denser 10%T gel^42^.

**Figure 2:**
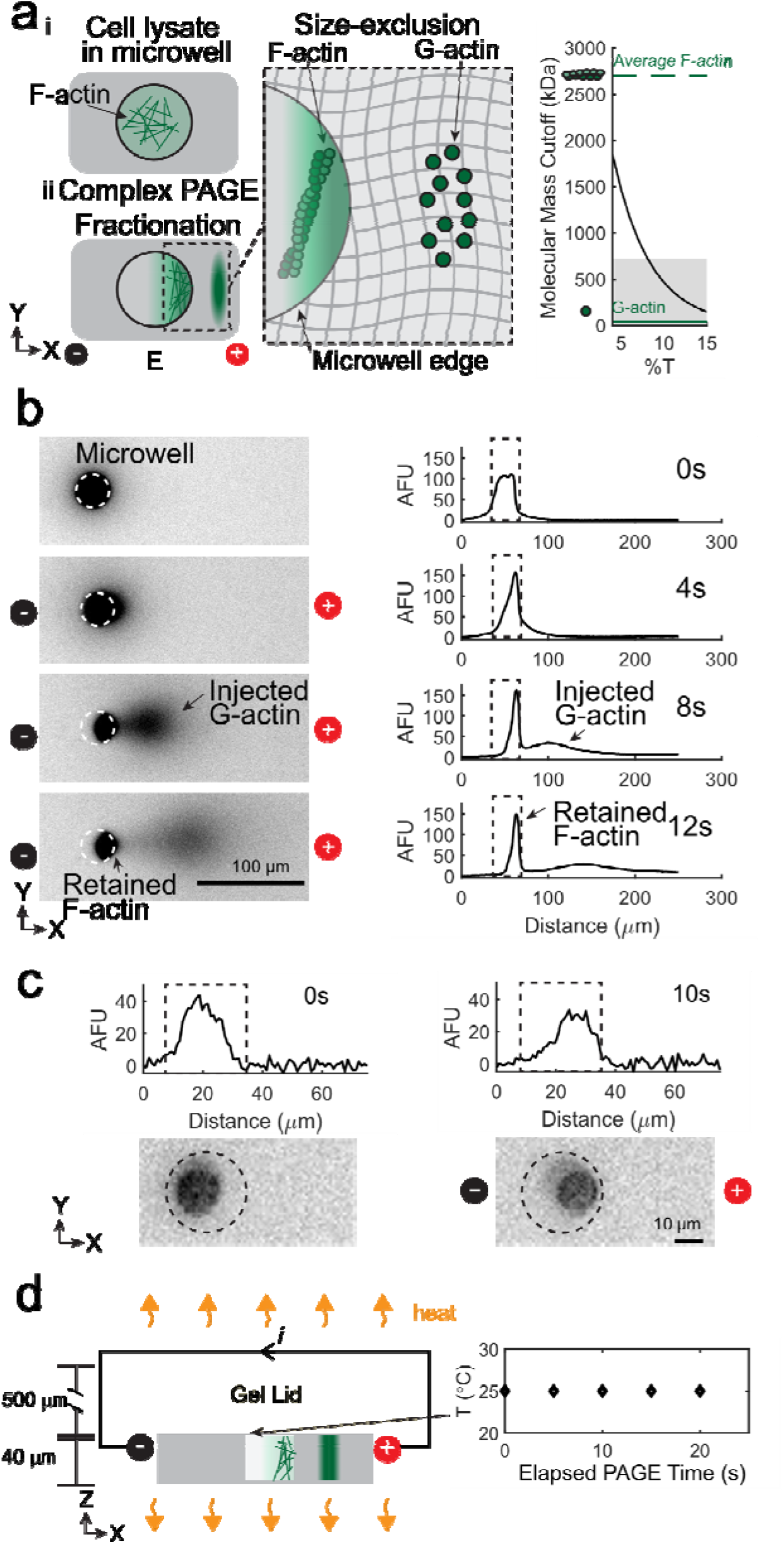
Size-based fractionation and efficient heat dissipation at the micro-scale provides molecular specificity to fractionate F-actin complexes from single cells. (a) Left: schematic of fractionation using polyacrylamide gel electrophoresis (PAGE) to separate F-actin complexes from G-actin monomers. Right: Estimated molecular mass cutoff as a function of gel density (%T). Shaded region is the molecular mass range of 99.9% of non-interacting protein species comprising the mammalian proteome, with notations indicating G-actin (42 kDa, solid green line) and average F-actin (~2700 kDa, dashed green line) molecular masses. (b) False-color fluorescence micrographs and corresponding intensity profiles during electrophoresis (30 V/cm) of MDA-MB-231 GFP-actin single-cell lysates in F-actin stabilization buffer; 76 ± 3% of the fluorescence remains in the microwell (n=4, error is standard deviation). (c) Intensity profiles (top) and false-color fluorescence micrographs of single RFP-Lifeact U2OS cells in microwells (dashed outline; only F-actin is fluorescent) upon lysis in F-actin stabilization buffer. PAGE results in retention of F-actin complexes in the microwell. (d) Left: schematic of heating in the fractionation gel (gray) and gel lid (yellow) upon applying a current. Right: plot of temperature as function of elapsed PAGE time under the F-actin stabilization lysis buffer gel lid at 30V/cm.

We further validate SIFTER maintains F-actin complexes during fractionation without PAGE-induced temperature rise that would depolymerize or dissociate protein complexes (e.g., above 45°C^43,44^). Electrical current passing through conductive buffer produces heat (Joule heating) during PAGE, which can increase temperature if not efficiently dissipated. The temperature difference, ΔT, between the surrounding medium and the conductor varies along the height axis, x, of the conductor: 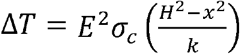, where *E* is the electric field strength (V/m), *σ_c_* is the buffer conductivity (S/m), *H* is the height and k is the thermal conductivity of the conductor (W/mK)^45^. Due to large temperature rises during electrophoresis in F-actin stabilization buffers containing MgCl_2_ (*σ_c_* ~1.3 mS/cm), *E* is limited to ~2-10 V/cm for 120-480 min in native slab gels^46^, or ~18 V/cm in capillaries^46^. In SIFTER, the anticipated ΔT at 30 V/cm is ~0.002°C (H~0.54 mm) *vs.* ~7°C increase in a slab gel (H ~ 5 mm; Supplementary Figure S2). Indeed, we measure constant room temperature using liquid crystal temperature sensors under the hydrogel lid during PAGE at 30 V/cm with SIFTER (Fig. 2d). Thus, we confirm SIFTER maintains endogenous protein complexes without Joule heating with ~100× faster fractionation than in a slab gel, 100-1000× higher sample throughput than a capillary (or comparable to automated capillary systems^47^), and without purifying, labeling or crosslinking of complexes^48^.

We sought to validate SIFTER’s quantification of single-cell heterogeneity of F-actin complex levels as quantitative assessment is needed for screening drugs targeting metastatic cell subpopulations^49^. In conventional imaging of F-actin with phalloidin (conjugated to a fluorophore), two factors pose a challenge to quantifying F-actin complex heterogeneity. First, phalloidin competes with or is dissociated from F-actin by both actin-binding proteins (e.g., cofilin)^19,20^ and drugs (e.g., actin nucleating drug jasplakinolide, Jpk^21^ and the structurally similar MiuA^50^). The number of potential actin-targeting drugs that compete with phalloidin are unknown. Nevertheless, Jpk and MiuA highlight the fact that a decrease in phalloidin staining signal can be due to decreased F-actin expression, competitive binding, or a combination of the two. Second, optimal cell segmentation requires that cells are not in contact with one another^51^, which limits quantification from tissues and high-throughput analysis^51^. The latter may be overcome in the case of actin by conducting analysis by flow cytometry. While flow cytometry is compatible with proximity ligation assay for two proximal proteins, the lack of antibodies specific for protein interactions prevents multicomponent protein complex measurement by flow cytometry^52^. Alternatively, SIFTER is free from competitive binding, cell segmentation challenges, and can discern and quantify protein complexes.

We investigated two well-studied F-actin drugs with SIFTER (Figure 3): Jpk and Latrunculin A (LatA)^53^. Understanding Jpk effects on F-actin complexes is confounded by competitive binding with phalloidin and differing observations *in vivo* versus *in vitro*^54^. Jpk lowers the number of actin subunits at which k_on_ becomes appreciable, causing disordered aggregates^54^. Still, F-actin complex levels increase in certain cell types with Jpk treatment in the 0.1 - 1.0 μM range as determined by bulk ultracentrifugation^55,56^. With phalloidin staining of Jpk-treated BJ fibroblasts, we qualitatively observe shorter filaments and small aggregates. When phalloidin stained Jpk-treated cells display decreased fluorescence signal, as with the BJ fibroblasts here, it is difficult to discern if competition with phalloidin obscures interpretation (Fig. 3a). SIFTER yields a ~1.7× and 2.7× decrease in median F-actin relative to the control at the 100 and 200 nM Jpk concentrations, respectively (Kruskal-Wallis p-value < 0.0001 with Dunn-Sidak correction for multiple comparisons, Fig. 3b-c). To assess heterogeneity in SIFTER F-actin complex levels across 100s-1000s of individual cells, we calculate the coefficient of quartile variation (CQV). The CQV is a metric of variance accounting for skewed distributions^57^, such as gamma-distributed protein expression^58^. The 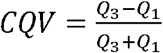, where Q_3_ is the 75^th^ percentile and Q_1_ is the 25^th^ percentile F-actin level. We find CQV_DMSO control, BJ_ =0.39, CQV_Jpk 100 nM, BJ_ =0.42 and CQV_Jpk 200 nM, BJ_ = 0.47 (subscripts refer to the treatment and cell type). Similar CQV values with increasing drug concentration indicate the drug effect is relatively consistent across the cell population.

**Figure 3:**
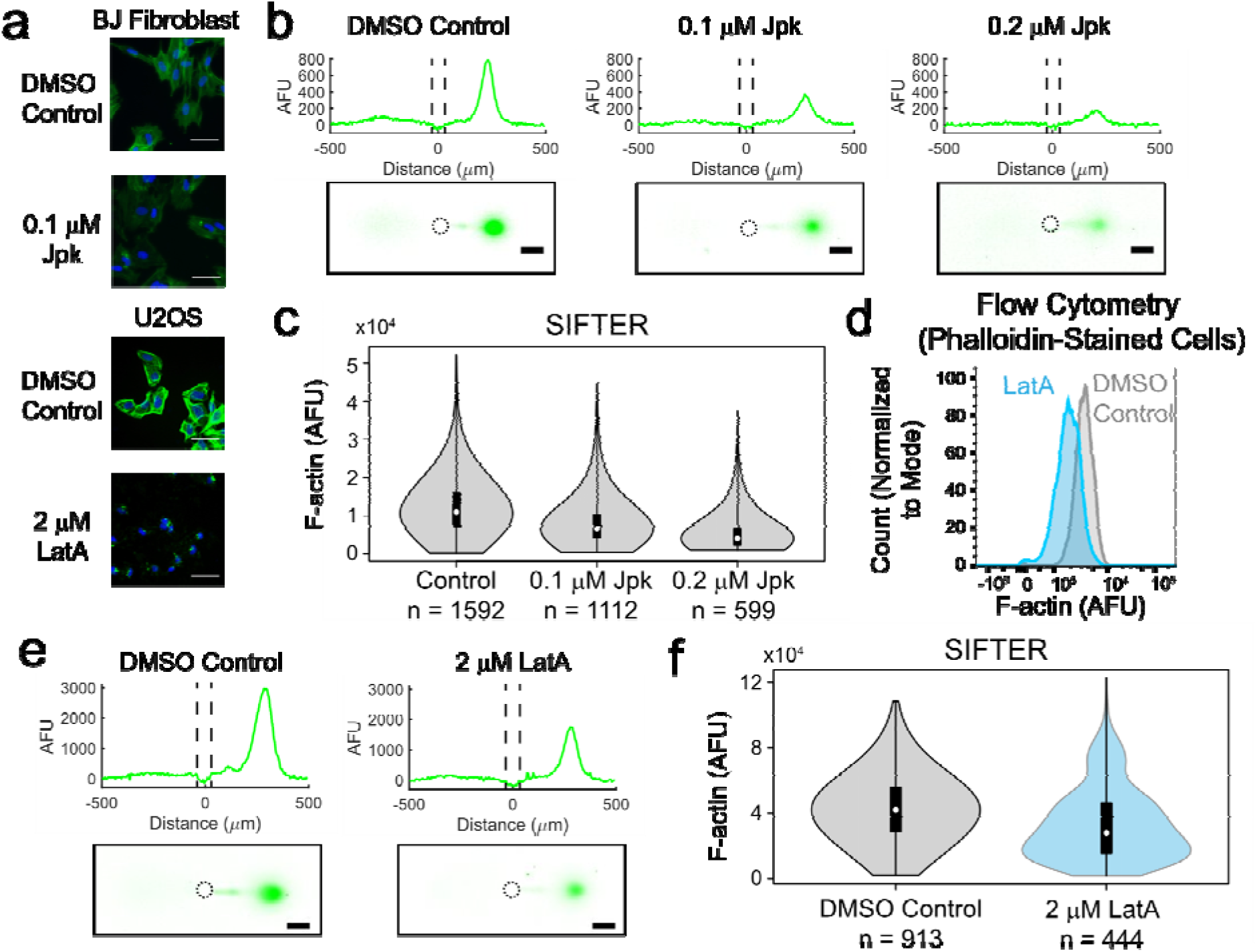
SIFTER quantifies cellular heterogeneity in F-actin complex levels, avoiding competitive binding or cell segmentation challenges encountered with phalloidin staining and capturing the cellular variation identified by flow cytometry. (a) False-color fluorescence micrographs of U2OS or BJ fibroblast cells fixed and stained with fluorescent phalloidin (F-actin, green) and Hoechst (nuclear stain, blue) after incubation with LatA (60 min) or Jpk (120 min). Scale bar is 50 μm. (b) False-color fluorescence micrographs and representative intensity profiles from SIFTER on single BJ fibroblast cells treated with the indicated concentration of Jpk. Scale bar is 100 μm. Microwell is outlined with a dashed line in the intensity profile (c) Violin plot of F-actin levels quantified from four different SIFTER devices with the indicated total number of single cells. Medians are 11026 for control, 6637 for 100 nM Jpk and 4041 for 200 nM Jpk. Kruskal-Wallis p-value < 0.0001 with Dunn-Sidak correction for multiple comparisons. (d) Histograms of F-actin fluorescence with cell count normalized to the mode from flow cytometry measurement of trypsinized and phalloidin-stained U2OS cells. Medians are 3454 for control (n = 9203) and 1858 for LatA (n = 5114). Mann-Whitney U Test p-value < 0.0001. (e) False-color fluorescence micrographs and representative intensity profiles from performing SIFTER on single U2OS cells. Scale bar is 100 μm. (f) Violin plot of F-actin levels quantified from four different SIFTER devices with the indicated total number of single cells. Medians are 42086 for control (n = 913) and 28144 (n = 444) for LatA. Mann-Whitney U test p-value < 0.0001.

LatA sequesters G-actin and reduces both F-actin complex levels and the *Fratio*, as determined by phalloidin staining and bulk methods, respectively ^51,57^ but variation in cell response is unknown. After treatment with LatA, we phalloidin stained U2OS cells and observe decreased F-actin complex-fluorescence (Fig. 3a) in agreement with previous findings^58^. To assess variation in cell response to LatA, we benchmarked the distribution of F-actin levels from LatA treatment in SIFTER versus flow cytometry of trypsinized, fixed, and phalloidin-stained U2OS cells. By flow cytometry, we find the median F-actin complex level of DMSO control cells is significantly higher than the LatA treatment median by 1.9× (Mann-Whitney P-value < 0.0001, Fig. 3d, n = 9203 control cells and n = 5114 LatA-treated cells). With SIFTER, we observe the median F-actin complex level in DMSO control cells is significantly higher than the LatA treatment median by 1.5× (Mann-Whitney P-value < 0.0001, n = 913 control cells, and n = 444 LatA-treated cells, Fig. 3d-e). We further found that SIFTER measured a significantly lower log fold change (i.e., smaller decrease in DMSO control over LatA) than flow cytometry (Mann-Whitney P-value < 0.0001, Supplementary Figure S3 and Supplementary Note 2). Thus, SIFTER does not measure as large a decrease in F-actin levels upon LatA treatment as flow cytometry of fixed and phalloidin-stained cells. One reason SIFTER may report smaller decreases in Factin levels upon LatA treatment (while still maintaining statistical significance) is due to run-to-run variation observed across assay replicates (each replicate shown in Supplementary Figure S4).

Interestingly, LatA treatment also corresponds with an increase in F-actin CQV as CQV_LatA, U2OS_ = 0.49 vs. CQV_DMSO control, U2OS_ = 0.32 by SIFTER (a 1.5× increase) and CQV_LatA U2OS_ = 0.30 vs. CQV_DMSO control, U2OS_ = 0.23 by flow cytometry (a 1.3× increase). Previously, phalloidin staining revealed a single F-actin complex-phenotype from ~200 sparsely seeded cells treated with 250 nM LatA^59^. Here, the CQV increase upon LatA exposure suggests differential cell tolerance to LatA potentially due to the almost 10× higher LatA concentrations utilized here. Thus, SIFTER circumvents competitive binding or cell segmentation challenges to quantify variation in drug effects on F-actin complexes at the single-cell level. The high CQV_LatA, U2OS_ from SIFTER prompted us to further investigate cellular variation in response to LatA treatment. It is not currently possible to quantify the variation in the other cytoskeletal protein complexes, IF and MT with flow cytometry, as vimentin and tubulin antibodies would bind both the monomer and protein complexes in the cell. However, with SIFTER, co-detection of protein complexes within the same cell is possible, using antibodies raised against different species, or with a chemical stripping and re-probing approach developed previously^31^.

### Multiplexed cytoskeletal protein-complex quantification uncovers compensation for actin perturbation in subpopulations of cells

We asked two questions regarding Latrunculin A-induced cellular variation, recognizing that SIFTER could permit measurement of all three major cytoskeletal protein complexes simultaneously. First, we sought to understand if LatA yields differential expression of other cytoskeletal protein complexes. Second, we asked whether LatA induced unique cell subpopulations. The cytoskeletal protein complexes F-actin, microtubules (MT, of α- and β-tubulin subunits), and intermediate filaments (IF, of vimentin or keratin subunits) have both redundant and distinct functions in maintaining cytoskeletal integrity (Figure 4a). Such redundancy^60^ yields increased IF to counteract F-actin destabilization of mesenchymal cells^61^ with another Latrunculin, LatB. Yet, quantification of cytoskeletal changes remains a challenge in single cells by microscopy due to segmentation artifacts and low signal-to-noise ratio from immunohistochemistry and phalloidin staining^62,63^.

**Figure 4:**
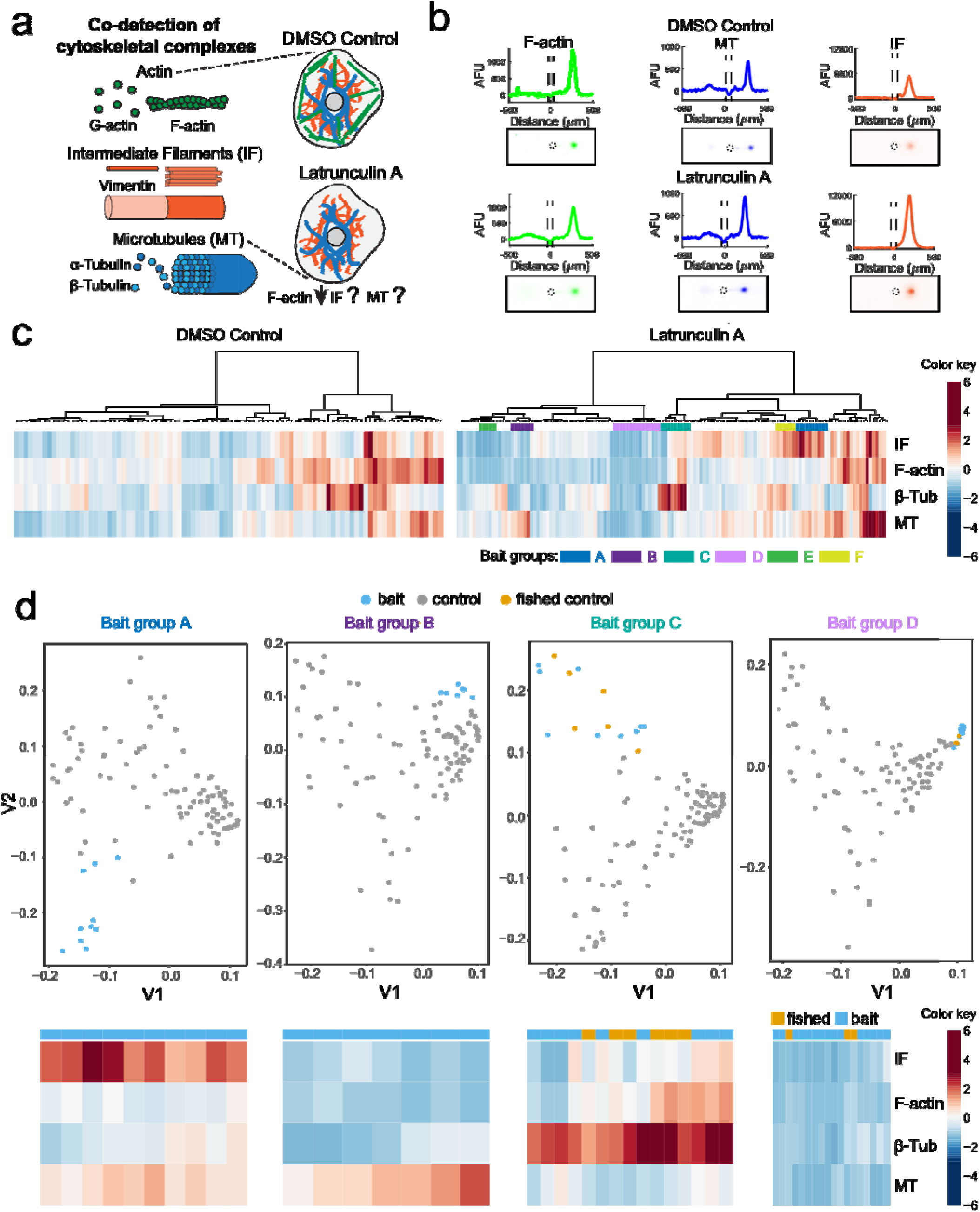
Multiplexed SIFTER detects subpopulations of cells with altered cytoskeletal protein complexes in response to F-actin destabilization. a) Schematic of the cell cytoskeleton composed of F-actin, intermediate filaments (IF) and microtubules (MT), and the unknown effects of Latrunculin A (LatA) on IF and MT. (b) Representative false-color fluorescence micrographs and intensity profiles from SIFTER. Monomeric proteins (e.g., β-Tubulin, β-Tub) are electrophoresed left of the microwell while F-actin, MT and IF are electrophoresed to the right of the microwell. Protein quantification is performed by peak area integration. Scale bar is 100 μm. (c) Heat maps with dendrograms from agglomerative hierarchical clustering with Euclidean distance metric and Ward linkage for U2OS cells incubated in DMSO (n = 92 cells, four SIFTER gels) or 2 μM Latrunculin A (LatA, n = 134 cells, four SIFTER gels). Distinct sub-lineages used as bait groups A-F for CellFishing are shown with colored bars. Heatmap is standardized by row (mean at 0, and color gradations at units of standard deviation). (d) Spectral clustering projections and heatmaps depicting LatA treatment bait group cells (blue), DMSO control cells (grey) and fished out DMSO control cells (yellow).

To understand concerted effects of LatA drug treatment on F-actin, MT, and IF, we performed same-cell, target-multiplexed SIFTER (Figure 4b and Supplementary Figure S5). We observe correlation between the three protein complexes in the DMSO vehicle control cells (n = 92 single cells), with Spearman rank correlation ρ = 0.66 for MT vs. F-actin, ρ = 0.64 for F-actin vs. IF, and ρ = 0.56 for MT vs. IF (Supplementary Figure S6; p < 0.01 for each correlation). While correlation suggests coordination of cytoskeletal protein-complex levels, agglomerative hierarchical clustering reveals cells with distinct patterns of protein-complex expression (e.g., groups A-F, Figure 4c).

Next, to elucidate whether any of the potential subpopulations shown in Figure 4c (e.g., groups A-F) were unique to the LatA-treatment, we adapted the GeneFishing method^64^ for “CellFishing”. Using a group of co-expressed cells as “bait”, we attempt to “fish out” other cells from a candidate pool that present a similar protein complex-expression pattern to that of the bait cells. We do this through a semisupervised clustering approach, coupled with sub-sampling to ensure robust discoveries. Here, groups of LatA-treated cells from hierarchical clustering that appear as unique phenotypes each define a set of “bait cells”, and the DMSO control cells define the candidate pool. If a group of bait cells does not identify any cells with similar phenotypes in the DMSO control cells, we assume the phenotype is unique to the LatA-treated cell population. We found that bait groups A, B, E and F do not fish out DMSO control cells, while groups C and D are examples of baits that do (Figure 4d). Group A (~11% of LatA-treated cells) is characterized by elevated IF and MT in response to F-actin destabilization. Note, 7 bait cells in Group A (Figure 4d) form a tighter and more distant sub-group from the DMSO control cells. Groups B and F (~8% and 7% of cells respectively) only sees the counteracting increase in either MT or IF, but not both. Compensation for F-actin perturbation by MT and/or IF in subpopulations of cells suggests these cells are better equipped to maintain cytoskeletal integrity in response to stress. Groups C (~10% of LatA cells) and D (~16% of LatA cells) both fish out small numbers of DMSO control cells (~6% and 3% of the DMSO control cells, respectively) and thus represent phenotypes not exclusive to LatA treatment. We hypothesize Groups D and E with low F-actin, MT, and IF may represent cells experiencing cytoskeletal collapse during apoptosis^65^ (LatA IC_50_ is ~0.5 - 3.0 μM with 24-hr exposure in breast and lung cancer cells)^66^.

### Quantifying distributions of total actin and *F_ratio_* across cells

To assess actin cellular heterogeneity, we asked: what are the statistical distributions of total actin and *F_ratio_* across cells? In order to assess statistical distributions across SIFTER replicates, we needed to measure the cells at a fixed time after preparing the single-cell suspension, as detachment lowers the level of cytoskeletal protein complexes^67-69^. We conducted SIFTER replicates with constant cell handling times and measured the *F_ratio_* from each device. The median *F_ratio_* values from the three SIFTER replicates were 0.48, 0.44 and 0.47 (n = 316, n = 253 and n = 123 respectively, Kruskal-Wallis p-value = 0.0152; Supplementary Figure S7), with a mean median of 0.46 and CV of mean median of 5%. The interquartile ranges were 0.14, 0.15, and 0.09, which indicates the distributions overlap substantially with similar medians despite statistically significant run-to-run variation indicated by the p-value < 0.05. In each of three replicates displayed as quantile-quantile (QQ) plots, we found total actin largely follows a gamma distribution, as expected based on transcriptional bursting (Supplementary Fig. S8)^70^. One of the replicates deviates from the gamma distribution at the highest quantiles, indicating the tail behavior is less well-described by a gamma distribution. For the first time, we find the *F_ratio_* follows a normal distribution across cells by examining the QQ plots (Supplementary Fig. S9). The normal *F_ratio_* distribution measured with SIFTER suggests actin binding proteins stochastically regulate actin polymerization/depolymerization.

Characterizing *F_ratio_* requires accurate quantification of the G-actin fraction to calculate the total actin (the denominator of the ratio, F+G). As with any immunoassay, immunoreagents must be screened for each specific application, as sample preparation determines whether the epitope is native, partially denatured, or fully denatured^71^ (Supplementary Table S1, Supplementary Note 3 and Supplementary Figure S8).

### Understanding cellular heterogeneity in actin distribution upon heat shock

To assess how a non-chemical stress perturbs (1) the *F_ratio_* distribution and (2) F- and G-actin coordination, we apply SIFTER to the study of heat shock. Cytoskeletal reorganization is a hallmark of disease states^5^, and protein-complex dysfunction is prominent in aging^72^ and during cellular stress^73,74^. Cell stresses such as heat shock yields re-organization of F-actin in many, but not all cell types^75^. Indeed, with phalloidin staining, we observed a qualitative decrease in F-actin fluorescence of RFP-Lentiviral transformed MDA-MB-231 GFP-actin cells upon heat shock (Fig 5a).

**Figure 5:**
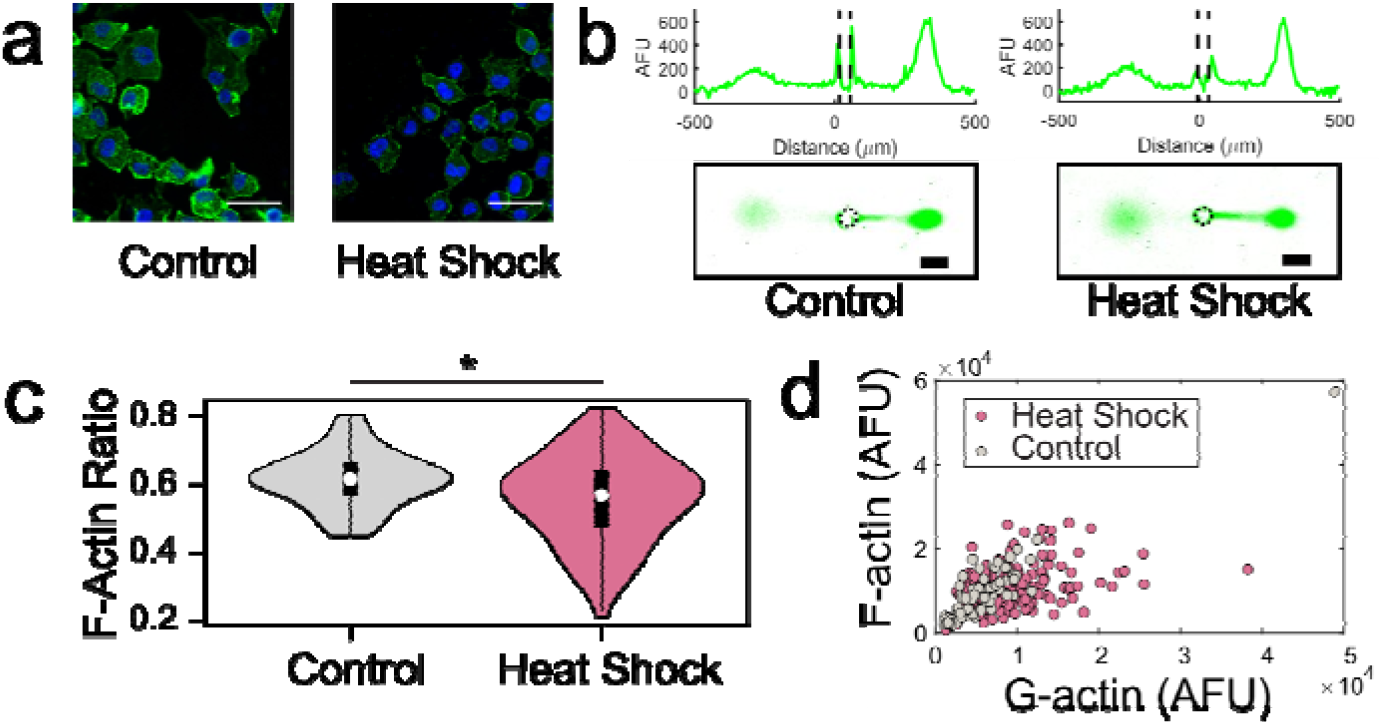
SIFTER quantifies actin distribution heterogeneity after heat-shock stress. (A) False-color fluorescence micrographs of adherent MDA-MB-231 GFP-actin cells with RFP-lentiviral transfection that were fixed and stained for F-actin (phalloidin, green) and the nucleus (Hoechst, blue) with heat shock (45°C for 60 min) or 37°C control. Scale bar is 50 μm. (B) Representative false-color fluorescence micrographs and intensity profiles of GFP-actin EP fractionation from the specified single cells. Scale bar is 100 μm. (C) Violin plots of F-actin ratio (F/F+G) from SIFTER with n = 81 for the control (one SIFTER gel) and n = 188 for the heat shock condition (two SIFTER gels). Median *F_ratio_* is 0.61 for control and 0.57 for heat shock. Mann-Whitney U = 12965 and the p-value is significant (*) at p = 0.0036. (D) Scatter plot of F versus G-actin. Spearman ρ = 0.82 for control and ρ = 0.42 for heat shock.

For more nuanced characterization of the *F_ratio_* distribution not possible with phalloidin staining, SIFTER reports the median *F_ratio_* in the heat-shocked cells was similar to control cells (0.57 *vs.* 0.61, respectively; Mann-Whitney p-value is 0.0036, Fig. 5b-c). However, the interquartile range of the *F_ratio_* in heat-shocked cells is nearly double that of control cells (0.16 *vs.* 0.09). We quantified the skew of the distribution with the Pearson’s moment coefficient of skew 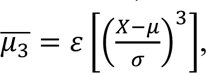 where ε is the expectation operator, X is the random variable (here, *F_ratio_*), μ is the distribution mean and σ is the standard deviation. We find *μ*_3_ is 0.08 for the control data set, and −0.32 for the heat-shocked cells with skew towards increased G-actin levels (Fig. 5d).

To understand if increased G-actin corresponds with discoordination of F- and G-actin levels upon heat shock, we quantified Spearman ρ (for F- and G-actin level correlation). The Spearman ρ decreased from 0.82 for the control to 0.42 for heat-shocked cells. Together, we conclude that F-actin levels alone cannot reveal cytoskeletal integrity: the *F_ratio_* distribution and Spearman ρ uncover differential stress response across the cell population.

## Discussion

SIFTER maintains multimeric cytoskeletal protein complexes during fractionation to reveal monomer versus protein-complex states in single cells. From perturbation of actin with well-characterized drugs, we find LatA, but not Jpk (at the concentrations tested), results in increased F-actin expression heterogeneity as characterized by increasing CQV. To investigate the heterogeneity of LatA-treated cells, we extended SIFTER to multiplexed readout of the three major cytoskeletal protein complexes (Factin, microtubules, and intermediate filaments) simultaneously in each cell. We identified previously unknown cell subpopulations, such as the cluster with decreased F-actin and compensatory increases in intermediate filaments and microtubules upon LatA treatment. Thus, observed heterogeneity in LatA Factin response corresponds with a spectrum of cytoskeletal integrity in the clonal population of U2OS cells investigated. While some cells increase expression of microtubules, intermediate filaments or both, other cells in the population undergo a complete cytoskeletal collapse. In the clonal population of U2OS osteosarcoma cells investigated here, the origins of differential maintenance of the cytoskeleton are unknown. However, recent single-cell sequencing studies of U2OS cells identified coordinated expression of sets of genes across subsets of cells, including some genes that regulate the cytoskeleton^76^. Partially coordinated regulation of the cytoskeleton raises two further questions: 1) what causes such differential gene expression to regulate the cytoskeleton in certain cells of a clonal population and 2) what is the functional implication of subsets of cells having a more resilient cytoskeleton? For the latter, we consider that the epithelial to mesenchymal transition of metastasis is marked by re-organization of the key cytoskeletal protein complexes^7^, and osteosarcoma is known for aggressive metastasis. Consequently, it is intriguing to consider whether cell subpopulations with compensating overexpression of microtubules and intermediate filaments (marked capability to re-organize the cytoskeleton) represent a more mesenchymal-like subtype.

Applying SIFTER to single-cell *F_ratio_* assessment, we determined, for the first time, that the F-actin ratio is normally distributed across a cell population. This indicates the possibility that the F-actin ratio could be a metric for assessing whether a population of cells is at an equilibrium state in terms of actin distribution. To investigate a non-chemical stress, we evaluated the impact of heat shock on the F-actin ratio of cells. We found marked F- and G-actin dysregulation and variability upon heat shock. Though missed by phalloidin staining, SIFTER uncovers marked skew in the F-actin ratio distribution upon heat shock. Our results present the possibility that SIFTER presents a more nuanced assessment of cytoskeletal integrity than phalloidin staining.

Cellular stresses, be they chemical, heat shock, hypoxia, or oxidative stress, are critical features of cancer biology. Understanding which protein complexes are differentially expressed in drug susceptible versus drug resistant cells, or in subsets of cells that metastasize will be critical to advancing cancer therapies. Thus, SIFTER unlocks the capability to assess single-cell heterogeneity in expression of multimeric protein complexes, with broad applications across biology, potentially including protein complexes unrelated to the cytoskeleton.

The SIFTER assay presently is conducted with a well-characterized F-actin stabilization buffer for cell lysis and maintenance of cytoskeletal protein complexes during electrophoresis. However, no single buffer is ideal for stabilization of all protein complexes, prompting careful optimization of detergent, salt (ionic species and concentration), buffer and pH for immunoprecipitation of specific sets of protein complexes^77^. We have not yet investigated alternative lysis buffers for SIFTER, such as certain immunoprecipitation buffers (e.g., containing 10-100 mM NaCl or KCl). Higher buffer salt concentrations than the F-actin stabilization buffer will increase buffer conductivity and we hypothesize could yield more extensive Joule heating that can dissociate protein complexes. Fabrication of thinner (<500 μm) hydrogel lids for efficient heat dissipation may be needed for PAGE in high salt buffers. Thus, further device or buffer optimization may be required to apply SIFTER to protein complexes beyond the cytoskeletal complexes investigated here.

The range of detectable and separatable protein-complex sizes is set by a tradeoff between fractionation and immunoprobing. Denser gels compromise assay detection sensitivity because size-exclusion based partitioning lowers the in-gel antibody probe concentration during the immunoassay^24^. Fractionation in decrosslinkable gel ^78^ should allow isolation of up to 100s of the known mammalian protein complexes with masses of ~295 kDa or greater in a 12%T gel (~7 or more protein subunits^79^, assuming each subunit has the average mammalian protein size of 375 amino acids^80^, or mass of ~40 kDa).

Another factor that determines which protein complexes are detectable with SIFTER is assay detection sensitivity. The cytoskeletal protein complexes investigated here are among the most abundant proteins in mammalian cells, often expressed at millions of copies^42^. Utilizing an in-gel immunoassay for readout, we have previously detected down to 27,000 copies of protein in a protein band^81^. As the SIFTER device is an open device design (vs. enclosed microchannels), protein is diffusively lost out of the microwell during cell lysis and out of the fractionation gel during electrophoresis. Such losses typically require proteins to be expressed at median copy number levels for mammalian proteins to be detectable in single-cell western blotting. While diffusive losses during SIFTER electrophoresis will be lower than in single-cell western blotting owing to efficient heat dissipation, protein fractionation inherently splits the amount of protein to be detected into the monomer and protein complex fractions. Thus, SIFTER likely requires proteins to be expressed above median copy numbers for detection.

One major advantage of SIFTER over existing assays for protein complexes^82^, such as FRET or proximity labeling, is that SIFTER measures endogenous proteins without requiring cell modifications. Thus, we anticipate SIFTER will be valuable in the measurement of protein complexes from clinical specimens. For example, our group previously introduced isolated circulating tumor cells into a microwell array single-cell western blot device for protein profiling ^83^. Circulating tumor cells are known to metastasize. With SIFTER, it would be informative to identify differentially expressed cytoskeletal protein complexes from circulating tumor cells to understand which protein complexes could be targets for small molecular inhibitors towards prevention of metastasis.

For time-sensitive cytoskeletal re-organization or mechano-sensitive protein complexes within the cytoskeleton (e.g., stress fibers and focal adhesions), the fractionation gel functionality can be extended to also serve as a cell culture extracellular matrix. On-chip culture can assay adherent cells without the perturbation of trypsinization^84^. We anticipate that SIFTER can aid in evaluating snapshots of dynamic processes while cells are still adherent, such as cytoskeletal recovery from acute stress (e.g., heat shock, hypoxia, etc.). In the present study, we trypsinized and gravity settled heat-shocked cells for 10 minutes after the heat shock stress. The amount of time for cytoskeletal recovery from heat shock depends on the duration of the heat shock and cell type, as mouse fibroblasts only partially restore F-actin within 24 hours after 1 hour at 43°C^75^. For shorter heat shock, or other stresses with faster recovery, growing and then stressing the cells on the SIFTER device will allow us to probe cytoskeletal protein-complex changes immediately after the stress, or at set times during the recovery. For mechano-sensitive cytoskeletal proteins, SIFTER may evaluate single-cell regulation of F-actin, MT, and IF in metastatic cancer cell subpopulations by quantifying dozens of cytoskeletal binding proteins with increased multiplexing by stripping and re-probing^81^. Looking ahead, SIFTER could assist drug screens targeting diverse protein interactions, and fundamental study of cellular stress responses underpinning invasive and heterogeneous cancer cells.

## Online Methods

### Chemicals

Tetramethylethylenediamine (TEMED, T9281), 40% T, 3.4% C acrylamide/bis-acrylamide (29:1) (A7802), N,N,N’,N’-, ammonium persulfate (APS, A3678), sodium deoxycholate (NaDOC, D6750), sodium dodecyl sulfate (SDS, L3771), bovine serum albumin (BSA, A7030), dithioerythritol (DTE, D8255), triton X-100 (X100), urea (U5378), β-Mercaptoethanol (M3148), anhydrous magnesium chloride (MgCl_2_, 814733) and dimethylsulfoxide (DMSO, D2438) were acquired from Sigma Aldrich. An Ultrapure Millipore filtration system provided deionized water (18.2 MΩ). PharmAgra Laboratories custom-synthesized N-[3-[(3-Benzoylphenyl)-formamido] propyl] methacrylamide (BPMAC). Phosphate buffered saline was purchased from VWR (10X PBS, 45001-130). Tris glycine (10X) buffer was obtained from Bio-Rad (25 mM Tris, pH 8.3; 192 mM glycine, #1610734). Petroleum jelly (Cumberland Swan Petroleum Jelly, cat. no. 18-999-1829). Tris-HCl was obtained from Fisher Scientific (1M, pH = 7.5; Corning MT46030CM), while 0.5 M Tris-HCl, pH 6.8 was purchased from Teknova (T1568). Photoinitiator 2,2-Azobis(2-methyl-N-(2-hydroxyethyl) propionamide) (VA-086) was acquired from FujiFilm Wako Pure Chemical Corporation. Gel Slick was purchased from Lonza (#50640). Tris Buffered Saline with Tween 20 (TBST-10X) was procured from Cell Signaling Technology (9997S). Paraformaldehyde (4% EM grade) was purchased from Electron Microscopy Sciences (157-4).

### Cell culture

All cell lines were authenticated by short tandem repeat profiling by the UC Berkeley Cell Culture facility and tested negative for mycoplasma. Naive U2OS cells were purchased from the UC Berkeley Cell Culture Facility. BJ fibroblasts were provided by the Dillin lab. U2OS RFP-Lifeact cells were previously generated by the Kumar lab^85^ at UC Berkeley, and kindly provided for this study. MDA-MB-231 GFP-actin cells were kindly provided by the Drubin lab at UC Berkeley. BJ fibroblasts and U2OS (RFP-Lifeact and naive) cells were maintained in DMEM (11965, ThermoFisher Scientific) supplemented with 10% FBS (100-106, GeminiBio), 1% penicillin/streptomycin (15140-122, ThermoFisher Scientific), and 1% non-essential amino acids (11140-050, ThermoFisher Scientific), while MDA-MB-231 GFP-actin cells were maintained in the same media minus the 1% non-essential amino acids. All cells were cultivated in a humidified incubator in 5% CO_2_ kept at 37 °C. Cells were sub-cultured at ~80% confluency and detached with 0.05% Trypsin-EDTA (Gibco #25300-054) for 3 min. Each SIFTER assay was performed on a distinct single-cell suspension.

### Generation of RFP-Lenti MDA-MB-231 GFP-Actin cells

MDM-MB-231 GFP-actin cells were a kind gift from the laboratory of Dr. David Drubin. Genome editing was performed at the genomic locus by integrating TagGFP (see Supplementary Methods) at the genomic locus for *ACTB*. Verification of genome editing was performed via standard PCR and sequencing. Briefly, DNA was collected from cells using the Qiagen DNeasy Blood and Tissue Kit (69506) as per manufacturer’s guidelines. 100 ng of genomic DNA was used for PCR using forward primer 5’GGACTCAAGGCGCTAACTGC3’ and reverse primer 5’ GGTACTTCAGGGTGAGGATGCC3’. Sequencing was performed using standard sanger sequencing using primer 5’GCTTCCTTTGTCCCCAATCTGG3’. A schematic for genome editing is provided in Fig. S10. MDA-MB-231 GFP-actin cells were infected with lentivirus containing CD510B-1_pCDH-CMV-MCS-ED1-Puro (SystemBio) modified to carry TagRFP (see Supplementary Methods and Supplementary Figure S10) under the CMV promoter.

### SIFTER assay

Buffers and gel lid incubation: F-actin stabilization lysis buffer used was 10 mM Tris-HCl, 1% Triton X-100, 2 mM MgCl_2_, and 0.5 mM DTE (titrated to pH=7.4). The DTE was added at the time of a given experiment. The depolymerization buffer was prepared as a 1.56x RIPA buffer such that upon addition of 8 M urea, the final buffer composition was 0.5x Tris glycine, 0.5% SDS, 0.25% sodium deoxycholate, 0.1% Triton X-100, 8 M urea, pH=8.3. Urea was added fresh at the time of the experiment and allowed to dissolve at 75°C. Hydrogel lids (15%T, 3.3% C) were photopolymerized as previously described between Gel Slick-coated glass plates offset with a 500 μm spacer^86^. Hydrogel lids were incubated overnight at 4°C in either the F-actin stabilization or the depolymerization buffer (before urea or DTE addition). Upon complete preparation of the urea-containing depolymerization buffer, the buffer was introduced to the gel lids in a water bath set to 75 °C and incubated for ~30 min before beginning the experiments. F-actin stabilization buffers and gel lids were kept at room temperature. Gel lids and buffers were only stored for up to 2 weeks, and buffer solution was never re-used.

Polyacrylamide fractionation gels (8%T and 3.3%C with 3 mM BPMAC incorporated) were polymerized on SU-8 micro-post molds as described elsewhere^31^. Trypsinized cells were introduced to the microwell array in 1X PBS solution for passive gravity settling. Trypsinization was performed for 3 min at 37°C, and cells in PBS (10010049, Thermo Fisher Scientific, pH=7.4, magnesium and calcium free) settled in the microwell array for 10 min. Each replicate experiment was run with a different 1-cm petri dish of freshly trypsinized cells in suspension.

For the fractionation separation, the fractionation gel device was pre-incubated in 10 mM Tris-HCl (pH=7.5) briefly before the glass slide was adhered to the surface of a custom 3D-printed PAGE chamber with petroleum jelly. A custom heater with a 12V PTC ceramic heating element (ELE147, Bolsen Tech) and PID temperature controller (ITC-106VH, Inkbird) was interfaced to the bottom surface of the PAGE chamber. The F-actin stabilization hydrogel lid was then applied to the array and cell lysis proceeded for 45 s before the electric field was applied (30 V/cm, 45 s for 42 kDa actin in U2OS or BJ fibroblasts, or 60 s for 69 kDa GFP-actin from the GFP-actin cells; Bio-Rad Powerpac basic power supply). Proteins were blotted, or bound to the fractionation gel, by UV-induced covalent immobilization to the BPMAC incorporated in the fractionation gel (Lightningcure LC5, Hamamatsu, 100% power, 45 s). The electrode terminals were reversed, and the hydrogel lid was exchanged with depolymerization buffer gel hydrogel lid for 45 s. PAGE was performed for the same duration in the opposite direction before a final UV photo-immobilization step (same UV power and duration). The glass slide was peeled from the PAGE chamber, and the fractionation gel was washed in 1X TBST for at least 30 min to overnight prior to immunoprobing.

Immunoprobing was performed as previously described^31^, utilizing a rabbit anti-GFP antibody for GFP-actin (Abcam Ab290), mouse anti-actin monoclonal antibody (Millipore MAB1501 for BJ fibroblasts), rabbit anti-actin polyclonal antibody (Cytoskeleton Inc. AAN01), rhodamine-labeled anti-actin Fab (Biorad 12004164 for BJ fibroblasts), rabbit anti-actin monoclonal antibody (Ab 200658 for BJ fibroblasts), rabbit anti-actin monoclonal antibody (Abcam Ab 218787 for U2OS cells), mouse anti-vimentin monoclonal antibody (Abcam Ab8978) and rabbit anti-β-tubulin monoclonal antibody (Abcam Ab6046). Gels were incubated with 50 μl of 1:10 dilution of the stock primary antibody in TBST for two hours and then washed 2x for 30 min in 1X TBST. Donkey Anti-Rabbit IgG (H+L) Cross-Adsorbed Secondary Antibody, Alexa Fluor 647-labeled (A31573, Thermo Fisher Scientific), Donkey Anti-Mouse IgG (H+L) Cross-Adsorbed Secondary Antibody, Alexa Fluor 555-labeled (A31570, Thermo Fisher Scientific) and Donkey Anti-Mouse IgG (H+L) Cross-Adsorbed Secondary Antibody, Alexa Fluor 647-labeled (A31571, Thermo Fisher Scientific) were used at a 1:20 dilution in TBST for a one-hour incubation after 5 min of centrifugation at 10,000 RCF. Two more 30-min TBST washes were performed prior to drying the gels in a nitrogen stream and imaging with a laser microarray scanner (Genepix 4300A, Molecular Devices). When immunoprobing with rhodamine-labeled anti-actin Fab and Ab 200658, 1:5 dilutions were used. For the Fab, immunoprobing completed after the two-hour incubation and two 30-minute washes in TBST. For multiplexed analysis of actin, vimentin and β-tubulin protein complexes, actin and vimentin were immunoprobed together, the gels were chemically stripped^31^ and then re-probed for β-tubulin. Chemical stripping was performed for at least one hour at 55°C. Gels were briefly rinsed in fresh 1x TBST three times and then washed in 1x TBST for at least one hour prior to re-probing.

Images were analyzed as described elsewhere^31^. Briefly, the images were median filtered utilizing the “Remove Outliers” macro in Fiji (pixel radius=2 and threshold=50 AFU). The images were then segmented, intensity profiles were generated for each separation lane by was fit to a Gaussian curve. For fits with an R^2^>0.7 and peaks with an SNR>3, user-based quality control is performed, and area under the curve is calculated within two peak widths from the center on the background subtracted profile. Image analysis was performed in MATLAB R2019b.

### Temperature measurement in SIFTER

Temperature sensors (liquid crystal thermometers; Type C 30-60 °C with 5 °C intervals from ThermometerSite) were placed directly under the hydrogel lid (immersed in F-actin stabilization lysis buffer). The temperature was monitored while applying 30 V/cm across the electrodes of the electrophoresis chamber.

### Fluorescence imaging of cells in microwells, lysis and PAGE

Imaging was performed via time-lapse epi-fluorescence microscopy on an Olympus IX50 inverted epifluorescence microscope. The microscope was controlled using Metamorph software (Molecular Devices) and images were recorded with a CCD camera (Photometrics Coolsnap HQ2). The imaging setup included a motorized stage (ASI), a mercury arc lamp (X-cite, Lumen Dynamics) and an XF100-3 filter for GFP (Omega Optical) and an XF111-2 filter for RFP (Omega Optical). Imaging was performed with a 10× magnification objective (Olympus UPlanFLN, NA 0.45) and 900 ms exposures with 1s intervals with U2OS RFP-Lifeact, and 2s exposure with 2s intervals with MDA-MB-231 GFP-actin (1x pixel binning). Exposure times were lowered for lysis imaging to 600 ms.

### F-actin cell staining with phalloidin and Latrunculin A and Jasplakinolide drug treatment

Latrunculin A (Cayman Chemicals 10010630) was dissolved in DMSO as a 2 mM stock solution and stored at −20 °C until use. Jasplakinolide (Millipore-Sigma, 420107) was reconstituted in DMSO and stored at −20 °C for up to 3 months. Cells were incubated in the drug solution at the concentration and for the time listed in the main text. The DMSO control cells were exposed to 0.1% DMSO in cell culture media for the same time as the drug treated cells. Cells were fixed with 3.7% paraformaldehyde in 1X PBS (10 min at room temperature), and permeabilized with 0.1% Triton X-100 (for 5 min at room temperature and stained with Alexa Fluor 647-labeled phalloidin (20 min at room temperature, ThermoFisher Scientific, A22287).

### Flow cytometry analysis of phalloidin-stained cells

Fixed cells were incubated in permeabilization buffer (0.1% Triton X-100 in PBS) at room temperature for 10 minutes. Cells were then spun down and incubated in staining solution (66 nM AlexaFluor 594 phalloidin in PBS supplemented with 2% BSA) at 4°C for 30 minutes. Finally, cells were washed twice with PBS and analyzed with flow cytometry using BD LSRFortessa.To analyze stained cells, single cells were gated by forward and side scatter. Only single cells were included in the fluorescence analysis.

### Heat shock treatment of cells

MDA-MB-231 GFP-actin RFP-lenti cells were incubated at 45 °C (VWR mini incubator, 10055-006) for 1-hour prior trypsinization and gravity settling in the fractionation gel.

### Statistical analysis

Mann-Whitney test (with U test statistic) and Kruskal-Wallis test with post-hoc Dunn’s test (Chi-squared test statistic), Spearman rank correlations, and QQ-plot generation with normal and gamma distributions were performed using pre-existing functions in MATLAB 2019b. All tests were two-sided. All boxplots include a centerline for the median, boxes at the 25^th^ and 75^th^ percentile and whiskers that extend to the extremes of the data. Violin plots were generated in RStudio (Version 0.99.903) using the library “Vioplot”. The boxplot within the kernel density plot displays boxes at the 25^th^ and 75^th^ percentile, a point at the median, and whiskers that extend to the extremes of the data.

### Cell Fishing clustering analysis

Standardization is by row for both the LatA treated and DMSO control data sets (expression level, or Gaussian protein peak AUC, for each protein complex) with the mean at 0 and standard deviation of 1. Initial agglomerative hierarchical clustering was performed separately for the LatA treated and DMSO control data sets utilizing Euclidean distances, and the Ward linkage criterion (R version 3.6.1, NMF package / MATLAB 2019b, Statistics and Machine Learning Toolbox). Distinct sub-clusters in the LatA treated data were further inspected as “bait” groups of cells inspired by the GeneFishing method described elsewhere^64^. We conducted an analogous analysis to GeneFishing, which we call “Cell Fishing”. Candidate cells from the DMSO control data sets were randomly split into subsamples of 23-33 cells, and each subsample was pooled together with the “bait” cells to form a sub-dataset. Semi-supervised clustering is applied to each sub-dataset using spectral analysis and a clustering algorithm based on the EM-fitted mixture Gaussian of two components model^87^ (R version 3.6.1, mclust package). The subsampling protocol was repeated 3000 times for a given “bait” set, and cells were considered “fished out” if they had a capture frequency rate of 0.99 or higher, as what is done in the GeneFishing paper^64^.

## Acknowledgements

This work was supported in part by the Office of the Assistant Secretary of Defense for Health Affairs under Award No. W81XWH-16-1-0002 (PI: Herr). The authors share their own conclusions and interpretation, which are not inherently endorsed by the Department of Defense. Other support includes: NSF Graduate Fellowship DGE1106400 (J.V.), Society of Lab Automation and Screening Graduate Education Fellowship (J.V.), NIH K99 (1K99AG065200-01A1, R.H.-S.), NSF CAREER CBET1056035 (A.E.H.), NIH R01CA203018 (A.E.H.), NIH R01AG055891-01 (A.D.) and Howard Hughes Medical Institute (A.D.). We thank the laboratories of David Drubin and Sanjay Kumar at UC Berkeley for providing the GFP-actin MDA-MB-231 and U2OS RFP-Lifeact edited cells, respectively.

## Availability of code

MATLAB functions to analyze protein expression levels from SIFTER gel images are available on Github (https://github.com/herrlabucb/summit). The GeneFishing algorithm for Python is also available on Github (https://github.com/tomwhoooo/GeneFishingPy).

## Availability of data

The data generated for this study are available in the Figshare repository with DOI: https://doi.org/10.6084/m9.figshare.c.5115779.v3

## Author Contributions

J.V., L.L.H, R.H.-S., C.K.T. and A.E.H. designed the experiments. J.V. and L.L.H. performed SIFTER assay and analysis, and phalloidin staining of adherent cells. C.K.T. provided cell culture and cell line modification support and conducted cell staining of suspension cells and flow cytometry. Y.Z. and H.H. designed and performed Cell Fishing clustering analysis and normalization and subsampling to compare flow cytometry and SIFTER F-actin distributions. All authors wrote the manuscript.

