## Supplementary Information for "Measuring expression heterogeneity of single-cell cytoskeletal protein complexes"

**Supplementary methods**

**Supplementary Figure S1:** Scatter plot and linear fit of protein molecular mass cutoff as a function of gel %T.

**Supplementary Figure S2:** Schematic representation of SIFTER and slab setup annotated with geometric parameters used for temperature model.

**Supplementary Figure S3:** Violin plots of the normalized log fold-change distributions of F-actin levels measured with flow cytometry and SIFTER.

**Supplementary Figure S4:** Boxplot of replicate DMSO control and LatA-treated U2OS cell measurements of F-actin by SIFTER from main text Figure 3F.

**Supplementary Figure S5:** Violin plots of microtubule and intermediate filament expression levels from single cells detected in main text Figure 4.

**Supplementary Figure S6:** Correlation matrix for actin filament, microtubule and intermediate filament expression levels from single cells detected in main text Figure 4.

**Supplementary Figure S7**: Boxplots of replicate SIFTER assays performed at a set time post-trypsinization.

**Supplementary Figure S8:** Quantile-quantile plots of total actin and F-actin ratio single-cell distributions.

**Supplementary Figure S9:** Annotated false color fluorescence micrograph of a SIFTER device immunoprobed with a gasket fixture to test multiple antibodies in distinct regions of the gel.

**Supplementary Figure S10:** Schematic of MDA-MB-231 genome edit with GFP.

**Supplementary Table S1:** Summary of immunoprobing results with various antibodies in SIFTER.

**Supplementary Note 1:** SIFTER buffer formulation and applicability to other cytoskeletal protein complexes.

**Supplementary Note 2:** Statistical analysis of flow cytometry and SIFTER distributions of F-actin levels in LatA versus DMSO control groups.

**Supplementary Note 3:** SIFTER immunoreagent screening.

**Supplementary methods:**
The tags inserted into the genome of the MDA-MB-231 cells were as follows:

TagGFP:
ATGGGATCCGGGGGCGAGGAGCTGTTCGCCGGCATCGTGCCCGTGCTGATCGAGCTGGACG
GCGACGTGCACGGCCACAAGTTCAGCGTGCGCGGCGAGGGCGAGGGCGACGCCGACTACG
GCAAGCTGGAGATCAAGTTCATCTGCACCACCGGCAAGCTGCCCGTGCCCTGGCCCACCCT
GGTGACCACCCTCTGCTACGGCATCCAGTGCTTCGCCCGCTACCCCGAGCACATGAAGATG
AACGACTTCTTCAAGAGCGCCATGCCCGAGGGCTACATCCAGGAGCGCACCATCCAGTTCC
AGGACGACGGCAAGTACAAGACCCGCGGCGAGGTGAAGTTCGAGGGCGACACCCTGGTGA
ACCGCATCGAGCTGAAGGGCAAGGACTTCAAGGAGGACGGCAACATCCTGGGCCACAAGC
TGGAGTACAGCTTCAACAGCCACAACGTGTACATCCGCCCCGACAAGGCCAACAACGGCCT
GGAGGCTAACTTCAAGACCCGCCACAACATCGAGGGCGGCGGCGTGCAGCTGGCCGACCA
CTACCAGACCAACGTGCCCCTGGGCGACGGCCCCGTGCTGATCCCCATCAACCACTACCTG
AGCACTCAGACCAAGATCAGCAAGGACCGCAACGAGGCCCGCGACCACATGGTGCTCCTG
GAGTCCTTCAGCGCCTGCTGCCACACCCACGGCATGGACGAGCTGTACAGG

TagRFP:
ATGAGCGAGCTGATCAAGGAGAACATGCACATGAAGCTGTACATGGAGGGCACCGTGAAC
AACCACCACTTCAAGTGCACATCCGAGGGCGAAGGCAAGCCCTACGAGGGCACCCAGACC
ATGAAGATCAAGGTGGTCGAGGGCGGCCCTCTCCCCTTCGCCTTCGACATCCTGGCTACCA
GCTTCATGTACGGCAGCAAAGCCTTCATCAACCACACCCAGGGCATCCCCGACTTCTTTAA
GCAGTCCTTCCCTGAGGGCTTCACATGGGAGAGAATCACCACATACGAAGACGGGGGCGT
GCTGACCGCTACCCAGGACACCAGCTTCCAGAACGGCTGCATCATCTACAACGTCAAGATC
AACGGGGTGAACTTCCCATCCAACGGCCCTGTGATGCAGAAGAAAACACGCGGCTGGGAG
GCCAACACCGAGATGCTGTACCCCGCTGACGGCGGCCTGAGAGGCCACAGCCAGATGGCC
CTGAAGCTCGTGGGCGGGGGCTACCTGCACTGCTCCTTCAAGACCACATACAGATCCAAGA
AACCCGCTAAGAACCTCAAGATGCCCGGCTTCCACTTCGTGGACCACAGACTGGAAAGAAT

CAAGGAGGCCGACAAAGAGACCTACGTCGAGCAGCACGAGATGGCTGTGGCCAAGTACTG
CGACCTCCCTAGCAAACTGGGGCACAGATGA

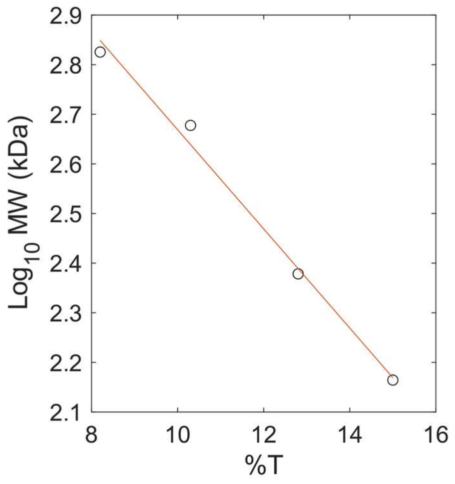


**Figure S1:** Scatter plot of protein size excluded in blue native PAGE as a function of Total acrylamide concentration (%T, g/mL) as reported by Wittig et al.[1] The linear fit is shown as a red line with equation: y = -0.0999T + 3.6683 ($r^2$ = 0.992).

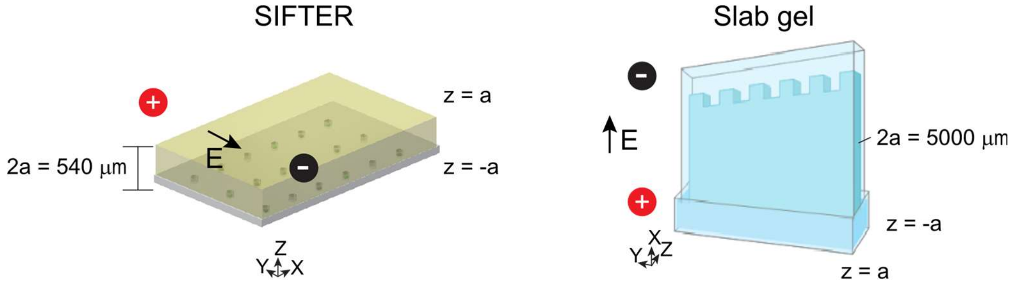


**Figure S2:** Schematic representations of SIFTER and a slab gel setup with parameters used for estimates of temperature difference. Temperature difference between the edge of the conductor and different z positions is given by[2]: $\Delta T = E^2 \sigma_c \left(\frac{a^2 - z^2}{2k}\right)$ where E is the electric field (V/m), $\sigma_c$ is the electrical conductivity (S/m), a is the cross-sectional thickness, and k is the thermal conductivity (W $m^{-1}$ $K^{-1}$). We neglect an additional term in the equation that accounts for heat transfer to a material incasing the conductor (thus we assume the width of the encasing material is zero). For E =3000 V/m, $\sigma_c$ = 0.13 S/m, and approximating the conductor as water (given the high water volume fraction of polyacrylamide gel), k = 0.5918 we find $\Delta T$=0.002°K in SIFTER (at the fractionation gel or z = -0.00004), and $\Delta T$ = 6.178 °K in the slab gel (at the location of the sample in the slab gel, or z = 0).

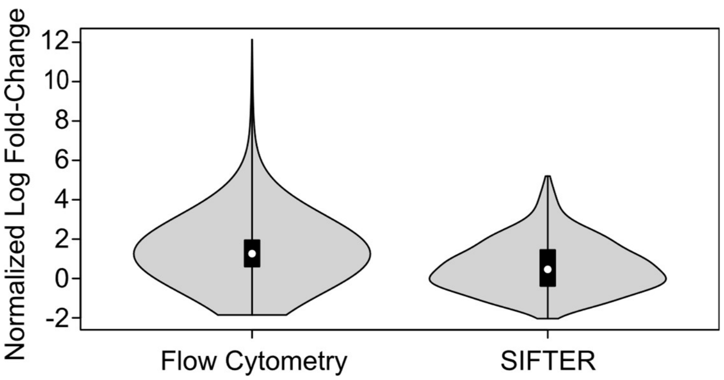


**Figure S3:** Violin plot of the normalized log fold-change distributions in F-actin levels from flow cytometry (of trypsinized, fixed and phalloidin-stained U2OS cells) and SIFTER from main text Figure 3. Normalization of DMSO control data to LatA data and subsampling results in a distribution with 44000 data points for flow cytometry and 444 data points for SIFTER, as described in Supplementary Note 2. Mann-Whitney p-value = 0 and the 99% confidence interval for a shift in locations is [0.6594, 0.9565].

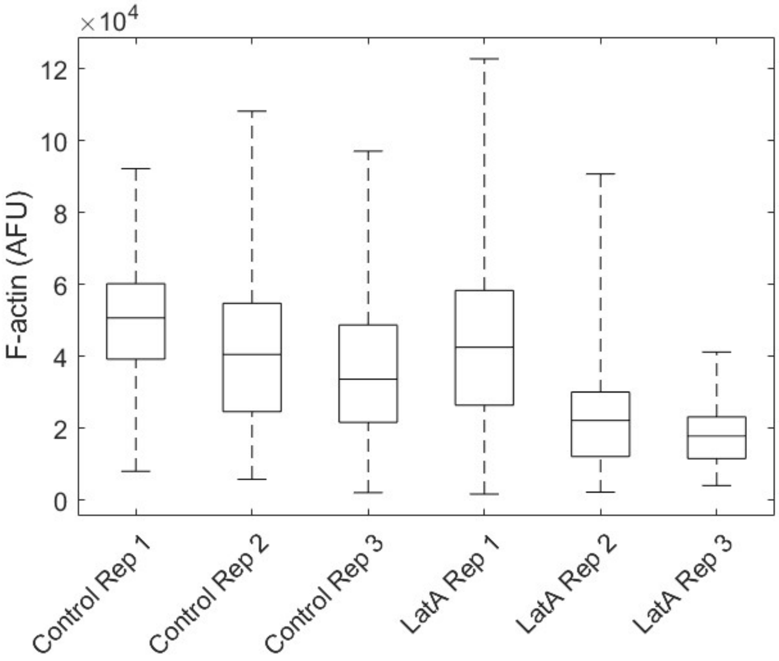


**Figure S4:** Boxplot of F-actin quantified from DMSO Control and LatA replicates (Rep) comprising main text Figure 3F. Kruskal-Wallis p-value $< 0.05$ for Control Rep 1-3 (n = 286, n = 288 and n = 339, respectively), and for LatA Rep 1-3 (n = 237, n = 97 and n = 110, respectively). Medians for Control Rep 1-3 are: 50749, 40599, and 33707, respectively. The DMSO Control mean median is 41685, and mean median CV = 21%. Medians for LatA Rep 1-3 are: 42581, 22255 and 17885, respectively. The LatA mean median is 27573 and mean median CV = 48%.

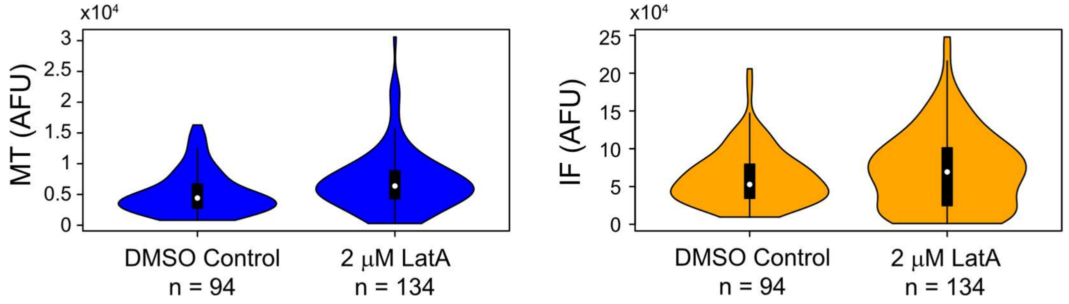


**Figure S5:** Violin plots of microtubule (MT) and intermediate filament (IF) expression levels with DMSO control or 2 μM LatA treatment. Mann-Whitney U = 8786 and p-values were: $p < 0.001$ for MT (DMSO control median = 4419 and LatA median = 6373). The Mann-Whitney U = 9945 with $p=0.309$ for IF (DMSO control median = 52754 and LatA median = 69349). Note: violin plots for F-actin are shown in main text Figure 3.

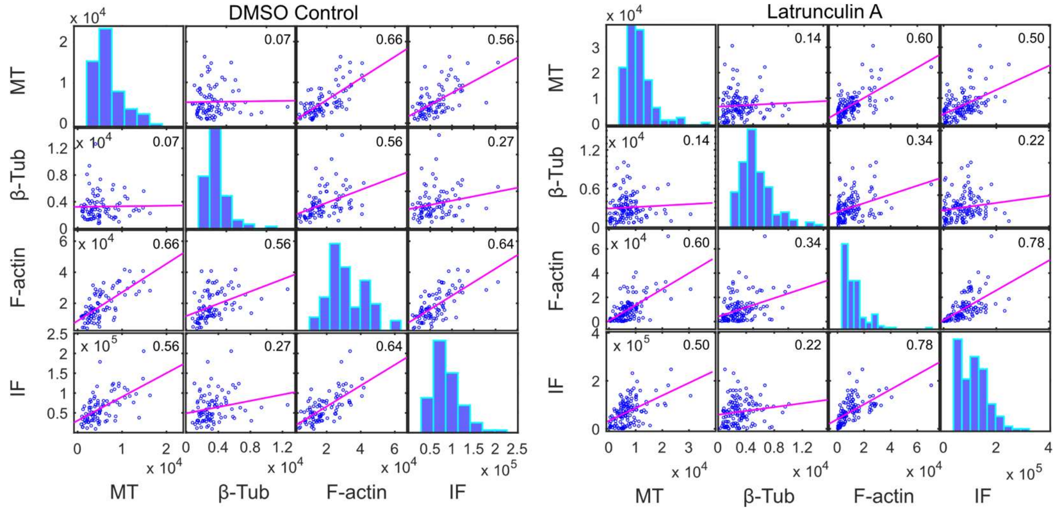


**Figure S6:** Correlation matrices for DMSO control and Latrunculin A treated cells. Protein complexes detected are microtubules (MT), F-actin and intermediate filaments (IF), and monomeric protein is β-tubulin (β-Tub). Spearman ρ are shown in the upper right-hand corner of each scatter plot (p-value < 0.01 for all correlations except B-tub vs. MT). Least-squares reference lines are shown in magenta with the slope equivalent to the correlation coefficient.

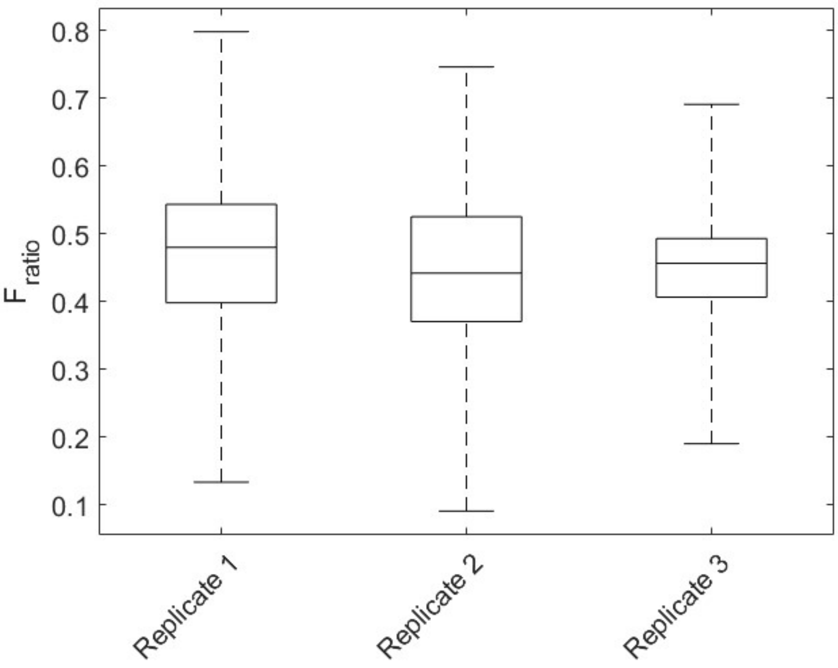


**Figure S7:** Boxplots of F-actin ratio for three SIFTER replicates performed on MDA-MB-231 GFP-actin cells (three different batches of cells with constant 10-minute settling time post-trypsinization performed on the same day). Replicate 1: n = 316; replicate 2: n = 253; replicate 3: n = 123. Kruskal-Wallis p-value = 0.0152; Dunn-Sidak post-hoc test for multiple comparisons p-values not significant except p = 0.0233 for Replicate 1 (median = 0.48) vs. Replicate 2 (median = 0.44).

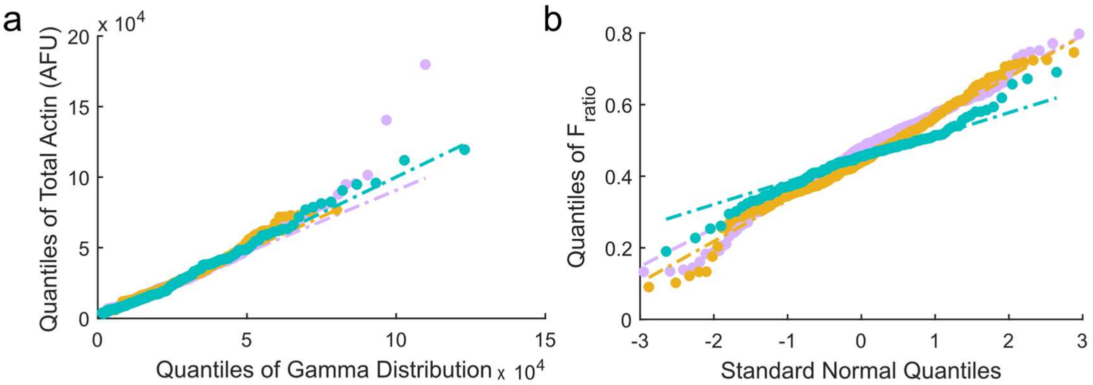


**Figure S8:** Quantile-Quantile (QQ) plots from replicates SIFTER assays in Fig. S3. (a) Total actin (F+G) for each single cell versus a gamma distribution to each replicate (replicate 1: purple; replicate 2: orange; replicate 3: green). Single-cell total actin for each single cell is indicated with a circle symbol and dashed lines represent the fitted gamma probability density function ($f(x;\alpha,\beta) = \frac{\beta^{\alpha} x^{\alpha-1} e^{-\beta}}{(\alpha-1)!}$ for each replicate. Fit parameter $\alpha$ = 3.5, 5.5 and 2.0, and $\beta$ = 9400, 5449, and 16108 for replicates 1, 2, and 3 respectively. (b) $F_{ratio}$ versus standard normal distribution.

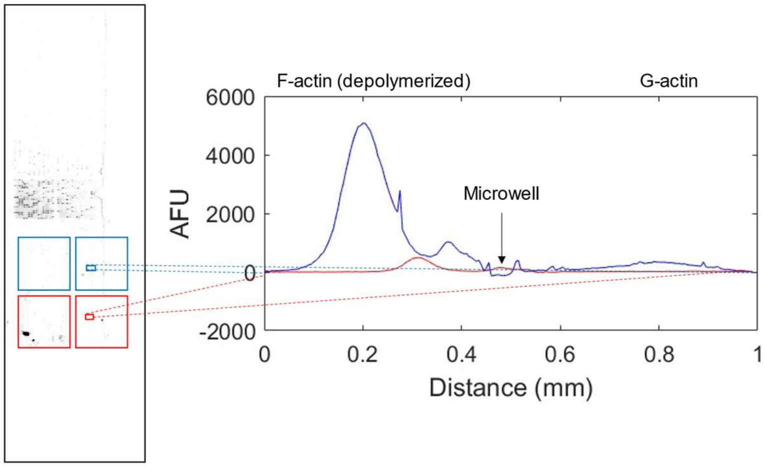


**Figure S9**: Gasket-based antibody screening for full SIFTER fractionation gels. Left: false-color fluorescence micrograph of fractionation gels (BJ fibroblast F and G actin separations) immunoprobed with different actin antibodies (Abcam Ab200658, blue; Abcam Ab198911, red).

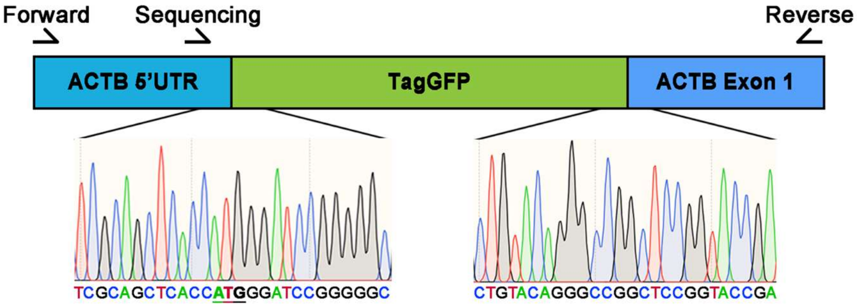


**Figure S10:** Schematic of genome edit generating the GFP-actin fusion in the MDA-MB-231 cells.

**Supplementary Table S1:** Summary of immunoprobing results with various antibodies.

| **Vendor** | **Product #** | **Clonality** | **Epitope Info** | **Valid Applications** | **Separation results** |
|---|---|---|---|---|---|
| Millipore | MAB1501 | Monoclonal (clone c4) | a.a. 50-70 (Chicken gizzard actin) | ELISA, IC, IF, IH, IH(P) & WB | F-actin band only (BJ fibroblasts) |
| CST | 8456S | Monoclonal | C-terminus of beta actin (synthetic peptide) | WB, IF, IHC | F-actin band only (BJ fibroblasts) |
| CST | 4968S | Polyclonal | Residues arounds Asp244 (synthetic peptide) | WB, IHC | F-actin band only (BJ fibroblasts) |
| Abcam | ab1801 | Polyclonal | ~residues 350-Cterminus (peptide) | WB, IHC | No signal (BJ fibroblasts) |
| Cytoskeleton Inc. | AAN01 | Polyclonal | synthetic peptide 11 C-terminal amino acids of actin | WB, ICC, ELISA | F-actin band only (BJ fibroblasts) |
| ThermoFisher | MA5-11869 | Monoclonal (clone c4) | Chicken gizzard actin | IF, IH(P), WB, IP | F-actin band only (BJ fibroblasts) |

| | | | | | |
|---|---|---|---|---|---|
| Abcam | ab198991 | Monoclonal | Synthetic peptide ~ amino acid 300 to C-terminus | WB, IP | F-actin band only (BJ fibroblast) |
| Abcam | ab200658 | Monoclonal | Synthetic peptide ~ amino acids 300 to C-terminus | WB, ICC, Flow | F-actin band only (BJ fibroblast and K562) |
| Abcam | ab218787 | Monoclonal | Synthetic peptide corresponding to human actin | ICC, WB | F-actin band only (U2Os and BJ fibroblast) |
| Biorad | 12004164 | Unspecified; rhodamine-labeled Fab | Recombinant human beta actin expressed in e. Coli | WB | F-actin band and some G-actin bands (BJ fibroblasts) |

**Supplementary Note 1:** We apply the F-actin stabilization buffer to the measurement of intermediate filament (IF) and microtubule (MT) cytoskeletal protein complexes. First, we note that MT and IF are relatively stable compared to F-actin. MT have depolymerization $t_{1/2}$ timescales of minutes[3] and IF experience subunit exchange ~10% over 7 hr[4]. Further due to the similarity in protein complex-stabilizing buffers for each cytoskeletal protein complex (Triton X-100 ~0.5-1.0%, pH ~6.7-7.4, and inclusion of 1 mM $MgCl_2$ for MT[5–7]), we determined the F-actin stabilization buffer employed in SIFTER could be usable for MT and IF fractionation.

**Supplementary Note 2:** For each of the two techniques, we converted the original data to the log scale, and normalized the LatA measurements by the mean and variance of the DMSO control. During normalization of the flow cytometry data set, sample size was accounted for by a repeated downsampling to match the sample size of SIFTER assay: we subsampled 444 points from 5114 flow cytometry LatA measurements and 913 points from 9203 flow cytometry DMSO control measurements. Next, we normalized the 444 subsampled LatA points, and repeated the subsampling and normalization protocol 100 times. The normalization procedure resulted in 444 SIFTER assay data points reflecting the log fold changes (DMSO control over LatA), and 44400 flow cytometry data points after pooling together the 100 subsamples. A Mann-Whitney test shows the normalized data in flow cytometry assay is significantly higher than for SIFTER assay, with a 0 p-value and 99% confidence interval of shift in locations being [0.6594, 0.9565]. This indicates that with a 99% high chance, the range [1.93, 2.60] will cover the ratio between the median fold change in flow cytometry and the median fold change in

SIFTER measurement (DMSO control over LatA). Here 1.93=exp(0.6594), 2.60=exp(0.9565). The confidence interval suggests the fold change measured in flow cytometry assay is significantly higher.

**Supplementary Note 3:** We found that in cells expressing endogenous (not GFP-fused) actin, some immunoreagents detected depolymerized F- but not G-actin (Supplementary Table S1, Supplementary Figure S5). Of note, a Fab fragment did yield G-actin immunoprobe signal in fibroblasts, while several full-length immunoreagents inconsistently detected G-actin in a subset of cells with F-actin signal. Lack of signal is not likely due to detection sensitivity, as actin is present at millions of copies of protein per cell[8] (while the in-gel immunoprobing limit-of-detection is ~27,000 copies of protein[9]). We instead hypothesize sterics may influence epitope availability. The Fab fragment may bind to native G-actin immobilized to the gel, whereas full-length antibody probes are unable to do so (~3× larger molecular mass). Future investigations will aim to establish a protocol for in-gel monomeric protein denaturation prior to immobilization to ensure the epitope availability is consistent between monomeric and depolymerized protein complex fractions immobilized in the gel.
